# Loss of TDP-43 function and rimmed vacuoles persist after T cell depletion in a xenograft model of sporadic inclusion body myositis

**DOI:** 10.1101/2021.04.09.439185

**Authors:** Kyla A. Britson, Jonathan P. Ling, Kerstin E. Braunstein, Janelle M. Montagne, Jenna M. Kastenschmidt, Andrew Wilson, Chiseko Ikenaga, William Tsao, Iago Pinal-Fernandez, Katelyn A. Russel, Nicole Reed, Kathryn R. Wagner, Lyle W. Ostrow, Andrea M. Corse, Andrew L. Mammen, S. Armando Villalta, H. Benjamin Larman, Philip C. Wong, Thomas E. Lloyd

## Abstract

Sporadic inclusion body myositis (IBM) is the most common acquired muscle disease in adults over age 50, yet it remains unclear whether the disease is primarily driven by T cell-mediated autoimmunity. IBM muscle biopsies exhibit nuclear clearance and cytoplasmic aggregation of TDP-43 in muscle cells, a pathologic finding observed initially in neurodegenerative disease, and nuclear loss of TDP-43 in neurons causes aberrant RNA splicing. Here, we show that loss of TDP-43 splicing repression, as determined by inclusion of cryptic exons, occurs in skeletal muscle of IBM patients. Out of 119 muscle biopsies tested, RT-PCR-mediated detection of cryptic exon expression is 84% sensitive and 99% specific for diagnosing IBM, indicating utility as a functional and diagnostic biomarker. To determine the role of T cells in pathogenesis, we generated a novel xenograft model by transplanting human IBM muscle into the hindlimb of immunodeficient mice. Xenografts from IBM patients display robust regeneration of human myofibers and recapitulate both inflammatory and degenerative features of the disease. Myofibers in IBM xenografts are invaded by human, oligoclonal CD8+ T cells and exhibit MHC-I upregulation, rimmed vacuoles, mitochondrial pathology, p62-positive inclusions, and nuclear clearance and cytoplasmic aggregation of TDP-43, resulting in expression of cryptic exons. Depletion of human T cells within IBM xenografts by treating mice intraperitoneally with anti-CD3 (OKT3) suppresses MHC-I upregulation, but rimmed vacuoles and loss of TDP-43 function persist. These data suggest that myofiber degeneration occurs independent of T cells, and muscle cell-intrinsic mechanisms, such as loss of TDP-43 splicing repression, drive IBM pathogenesis.

**One Sentence Summary:** Depletion of T cells in a xenograft model of sporadic inclusion body myositis suppresses inflammation but not TDP-43 pathology or muscle degeneration.

## Main Text

### INTRODUCTION

Sporadic inclusion body myositis (IBM) causes progressive muscle atrophy, weakness and disability, with a median of 14 years between symptom onset and wheelchair-dependence (*1–3*). The insidious progression of weakness and atrophy makes the disease difficult to diagnose, with a median time from onset to diagnosis of about 5 years (*4–6*). Clinically, IBM is suspected based on the pattern of muscle weakness with finger flexor and quadriceps muscles most severely affected. Diagnosis is confirmed by muscle biopsy, which shows characteristic features including ubiquitinated protein inclusions, rimmed vacuoles, mitochondrial pathology, and intense endomysial inflammation (*1*). Although many clinical trials of immunomodulatory drugs have been performed in IBM, as well as trials of drugs targeting proteostasis or the myostatin pathway, none have proven successful, and there remains no established therapy for the disease (*7*).

The combination of inflammatory and degenerative pathological features has led to considerable debate as to the primary cause of the disease (*7–10*). The presence of major histocompatibility complex class I (MHC-I) upregulation within myofibers that are invaded by highly differentiated cytotoxic CD8+ T cells, the association of IBM with other autoimmune disorders and specific Human Leukocyte Antigen (HLA) loci, and the presence of autoantibodies in IBM patient sera support an autoimmune trigger for the disease (*11–14*). However, the fact that immunosuppression fails to show clinical benefit suggests that endomysial inflammation may not be required for disease progression. Alternatively, IBM pathology may be driven by a T cell population that is refractory to immunosuppressive therapy (*11, 15*).

A competing hypothesis is that IBM is primarily a degenerative disease, analogous to neurodegenerative diseases. This view is supported by ultrastructural characterization of the inclusions in IBM which form amyloid-like fibrils and tubulofilaments analogous to those observed in the brain in Alzheimer’s disease (AD) (*16*). Indeed, intracellular β-amyloid accumulation is observed in IBM muscle, and overexpression of amyloid β precursor protein (APP) in human myoblasts or mouse muscle causes toxicity (*17, 18*). More recently, nuclear loss and cytoplasmic inclusions of transactivation response element DNA-binding protein 43 (TDP-43) have been observed in IBM patient myofibers (*19–21*), a pathological feature initially identified in patients with Amyotrophic Lateral Sclerosis (ALS) and Frontotemporal Dementia (FTD) (*22*). By binding to the UG-rich repeats of their target RNAs to repress splicing of nonconserved cryptic exons in a transcriptome-wide manner (*23–25*), TDP-43 serves to protect cells by ensuring the proper formation of the transcriptome (*26*). TDP-43 is recognized as the founding member of a growing family of RNA binding proteins that serve this critical cellular function (*27*). Recent studies in human disease support the view that loss of TDP-43 splicing repression underlies neuronal loss in neurodegenerative diseases (*26, 28, 29*). While a pathological link between IBM pathogenesis and age-dependent neurodegeneration exists, it remains unclear whether loss of TDP-43 splicing repression occurs in skeletal muscle of IBM patients.

A primary reason that pathogenesis of IBM remains elusive is that there are no laboratory models of IBM that recapitulate both the inflammatory and degenerative features of the disease. Patient-derived xenograft (PDX) models have been effectively utilized to develop laboratory models and treatments for malignancies, as well as personalized therapies (*30–35*). Xenografts have recently been used to study genetic disorders of skeletal muscle, and whole muscle xenografts have been successfully used to model fascioscapulohumeral muscular dystrophy and test potential therapies (*36–38*). In a novel application of this methodology, we aimed to determine whether human skeletal muscle xenografts could be used to model sporadic, late-onset, inflammatory muscle diseases, such as IBM. Aging is known to diminish the regenerative capacity of skeletal muscle, resulting in sarcopenia, the loss of muscle mass and strength observed in elderly populations (*39–41*). In addition, *in vitro* studies have shown myoblasts isolated from IBM patients proliferate at a slower rate than age-matched controls and show telomere shortening, indicative of premature senescence (*42*). Furthermore, skeletal muscle regeneration is highly regulated by the immune system, and the chronic inflammation in IBM has been hypothesized to negatively influence muscle regeneration (*43–46*). Conversely, it has been reported that IBM patient muscle exhibits increased numbers of satellite cells and regenerating fibers in comparison to age-matched controls, as well as upregulation of the myogenic regulatory factor myogenin, which drives differentiation of satellite cells (*47, 48*). Since xenograft formation requires successful myofiber regeneration, this approach allows us to test the ability of human IBM muscle to regenerate *in vivo* as well as develop a laboratory model for IBM.

Here, we show that the detection of cryptic exons in human muscle is a sensitive approach to identify TDP-43 pathology in IBM patient muscle, and these cryptic exons may have utility as biomarkers. Importantly, these cryptic exons can be detected in IBM xenografts, which regenerate normally in comparison to controls and recapitulate both inflammatory and degenerative pathological features of the human disease. We can use this model to carry out mechanistic studies to better understand disease pathogenesis. By treating mice containing IBM xenografts with a monoclonal CD3 antibody (OKT3), we can specifically deplete human T cells from xenografts in order to assess what pathological features of IBM are dependent on T cells. Our data suggest that while T cells are required for MHC-I upregulation and mitochondrial dysfunction, muscle cell-intrinsic mechanisms, such as rimmed vacuole formation and loss of TDP-43 function, may be the primary driver of muscle degeneration in this xenograft model of IBM.

### RESULTS

#### Detection of TDP-43 cryptic exons in muscle is a sensitive and specific assay for diagnosing IBM

Of the many proteins found to form aggregates within IBM patient muscle biopsies, cytoplasmic accumulations of TDP-43 and p62/SQSTM1 have been suggested to have high specificity for IBM (*19, 20*). In ALS, AD, and other neurodegenerative diseases termed TDP-43 proteinopathies (*49, 50*), nuclear loss of TDP-43 is more frequent than cytoplasmic aggregation, and assays for loss of TDP-43 function are more sensitive for detecting TDP-43 pathology than immunohistochemical stains for TDP-43 localization (*26, 51*). TDP-43 normally functions in mRNA splicing to repress the incorporation of “cryptic” exons, and detection of cryptic exons in neurodegenerative disease tissue using RNA-sequencing and RT-PCR has been shown to be a sensitive functional assay for TDP-43 pathology (*26, 52*). Indeed, cryptic exon incorporation can be detected in hippocampal samples from AD cases lacking TDP-43 aggregation (*51*). Whether loss of TDP-43 splicing repression occurs in skeletal muscle of IBM patients remains to be determined. Since nonconserved cryptic exons are species- and cell type-specific, we first sought to identify human skeletal muscle-associated TDP-43 cryptic exon targets (*26, 52*).

To identify TDP-43 cryptic exons from muscle cells, we used previously characterized siRNAs (*26*) to deplete TDP-43 from cultured, healthy human myoblasts (**Fig. 1A**). Total RNAs extracted from these TDP-43-deficient myoblasts were subjected to RNA-seq analysis, and we identified a set of RNA targets in which nonconserved cryptic exons were flanked by “UG” repeats, the known binding site of TDP-43 (**Fig. S1**). In addition to cassette exons, we identified a subset of non-standard cryptic exons that could be categorized as alternative transcriptional start sites, premature polyadenylation sites, or expansions of conserved canonical exons (**Fig. 1B**, **Table S1**). These results confirm that TDP-43 represses nonconserved cryptic exons in human myoblasts.

**Fig. 1.**
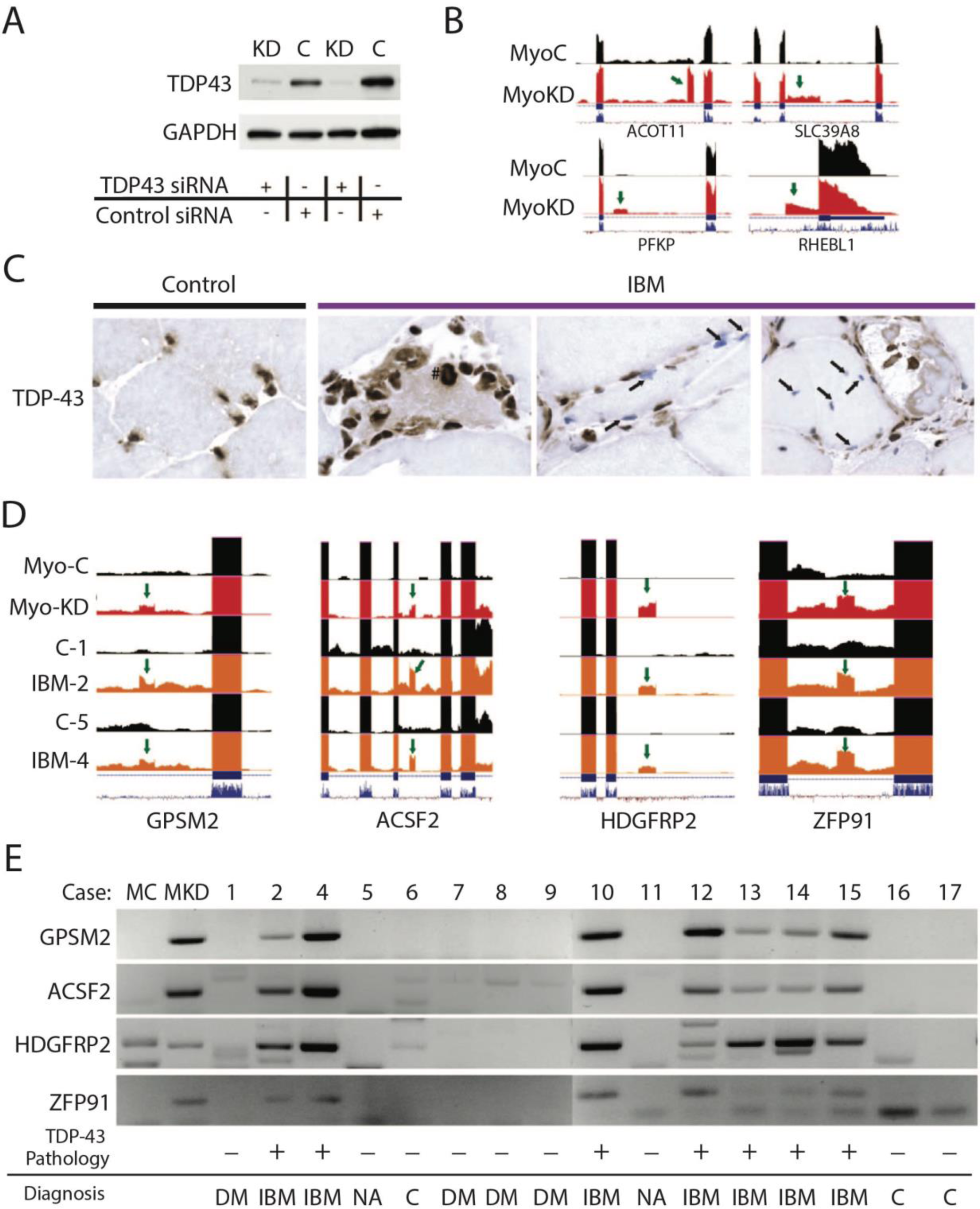
Cryptic Exon detection is a sensitive and specific assay for TDP-43 pathology in IBM biopsies. (A) Levels of TDP-43 are substantially reduced in myoblasts treated with TDP43 siRNA (KD) compared to control siRNA (C). (B) Visualization of the cryptic exons (green arrow) in myoblast cells with TDP-43 knockdown (MyoKD) compared to control myoblasts (MyoC): ACOT11, SLC39AB, PFKP, and RHEBL1. (C) Immunohistochemical TDP-43 staining of muscle sections: control muscle showing TDP43 staining in the nucleus, and IBM muscle showing accumulation of TDP43 in the cytoplasm (#) or nuclear clearing of TDP43 (arrows). (D) Visualization of the cryptic exons (green arrow) in myoblasts (MyoKD or MyoC) or patient muscle biopsies (control (C) and IBM; numbers indicate cases (Table S2)). (E) Representative gel showing cryptic exon expression from TDP-43 target genes GPSM2, ACSF2, HDGFRP2 and ZFP91 in skeletal muscle biopsies from IBM and control biopsies (DM: dermatomyositis; NA: neurogenic atrophy; C: normal muscle or mild nonspecific features).

If loss of TDP-43 function occurs in IBM, there should be an incorporation of these nonconserved cryptic exons in skeletal muscles of IBM patients. To directly assess whether repression of cryptic exons is compromised in IBM, we first confirmed the diagnosis from muscle biopsies of a cohort of cases that exhibit pathological hallmarks of IBM, including rimmed vacuoles, endomysial inflammation, and p62 and TDP-43-positive protein aggregates (**Table S2**). As shown previously (*53*), besides the abnormal cytoplasmic accumulation of TDP-43, we also found clearance of TDP-43 from nuclei in surrounding but otherwise morphologically normal myofibers (**Fig. 1C**).

Total RNA from biopsies of quadriceps muscle from two IBM patients along with two controls were extracted and subjected to RNA-sequencing analysis (**Table S2**). Importantly, we found several nonconserved cryptic exons in both IBM cases but not controls (**Fig. 1D**). To determine whether the incorporation of cryptic exons can be readily detected in IBM patient muscle, total RNA was extracted from rectus femoris muscle biopsies of age- and sex-matched ENMC clinically-defined IBM and control patients (n=16, **Table S2)** and subjected to RT-PCR analysis. We selected a panel of four RNA targets (GPSM2, ACSF2, HDGFRP2, and ZFP91) for our analysis, and sequence and alignment validation of cryptic exon RT-PCR products were performed for the RNA targets (**Fig. S2, S3**). Remarkably, cryptic exons of all four genes were detected in all 7 IBM cases, but not in any of the 9 controls (**Fig. 1E**, **Table S3**). Together, these results establish that loss of TDP-43 function occurs in IBM patient biopsies.

To determine the sensitivity and specificity of cryptic exon detection for the diagnosis of IBM in a large cohort of myositis center patients, we assayed muscle biopsies from an additional 103 patients (**Table S3**), including 30 patients used in generating the xenograft model (see below). Out of a total of 119 total patients, 36/44 IBM patients and only 1/75 control patients were positive for cryptic exons, resulting in a sensitivity of 84% and specificity of 99% for diagnosis of IBM. These data show that detection of TDP-43 cryptic exons represents a functional biomarker and is a sensitive and specific assay for IBM diagnosis among myositis patients.

#### IBM xenografts regenerate robustly in immunodeficient mice

To determine whether human skeletal muscle xenografts transplanted into immunodeficient mice can successfully engraft and recapitulate key features of human muscle pathology, we recruited subjects at the time of diagnostic open muscle biopsy in the Johns Hopkins Outpatient Center (**Table S4**). Patients were classified as sIBM (n = 15) if they met ENMC 2011 criteria or as a control (n = 12), if they were over 40 years of age and IBM was not considered to be a possible diagnosis at follow-up (*4, 54*). The IBM biopsies display characteristic features of the disease, including endomysial inflammation, invasion of myofibers by CD3+ T cells, cytochrome c oxidase (COX)-deficient fibers, and/or rimmed vacuoles (**Table S4**). The control biopsies were taken from patients with weakness, elevated muscle enzymes, and/or muscle pain, and typically showed mild abnormalities such as perivascular inflammation, mild necrosis, myophagocytosis, mild myopathic features (e.g., scattered internalized nuclei), mild neurogenic atrophy, and/or mitochondrial abnormalities, though 5 of 12 were pathologically normal. The sex ratio and disease duration of the patients are not significantly different between groups; however, the IBM patient population is older than the control group (average age 70.1 vs 61.2 years, p = 0.04) (**Table S5**).

Human skeletal muscle biopsy specimens obtained from both IBM and control patients were dissected and transplanted into NOD-Rag1^null^IL2rγ^null^ (NRG) mice lacking the ability to generate mature B or T cells and innate lymphoid cells, including NK cells (**Fig. S4A**) (*37, 55*). In this model, the mature human myofibers cut during the biopsy degenerate and are replaced by newly regenerated myofibers, derived from transplanted satellite cells, that are revascularized and reinnervated by the mouse host (*37, 55*). At 1 month after xenotransplantation, all myofibers in IBM and control xenografts are positive for embryonic myosin heavy chain (eMHC), a marker of newly regenerated myofibers (**Fig. S4B-C**) (*37, 55*). At 4 months of age, myofibers within IBM xenografts successfully regenerate as well as control xenografts (**Fig. 2A-E**). There are no significant differences in the number of regenerated fibers or the fraction of the xenograft comprised of myofibers (fiber fraction) between the two groups (**Fig. 2B,C**), and the percent of embryonic myosin heavy chain (eMHC) positive fibers is unchanged (**Fig. 2D**), indicating that the process of myofiber maturation, as assessed by the turnover of eMHC, is unaltered (*56*). Interestingly, myofibers of IBM xenografts show a significant increase in the median cross-sectional area (CSA) in comparison to control xenografts (395 vs. 249 μm^2^, p = 0.011) (**Fig. 2E**). These data indicate IBM patient muscle is capable of robust regeneration within a mouse host, allowing the establishment of a novel IBM model.

**Fig. 2.**
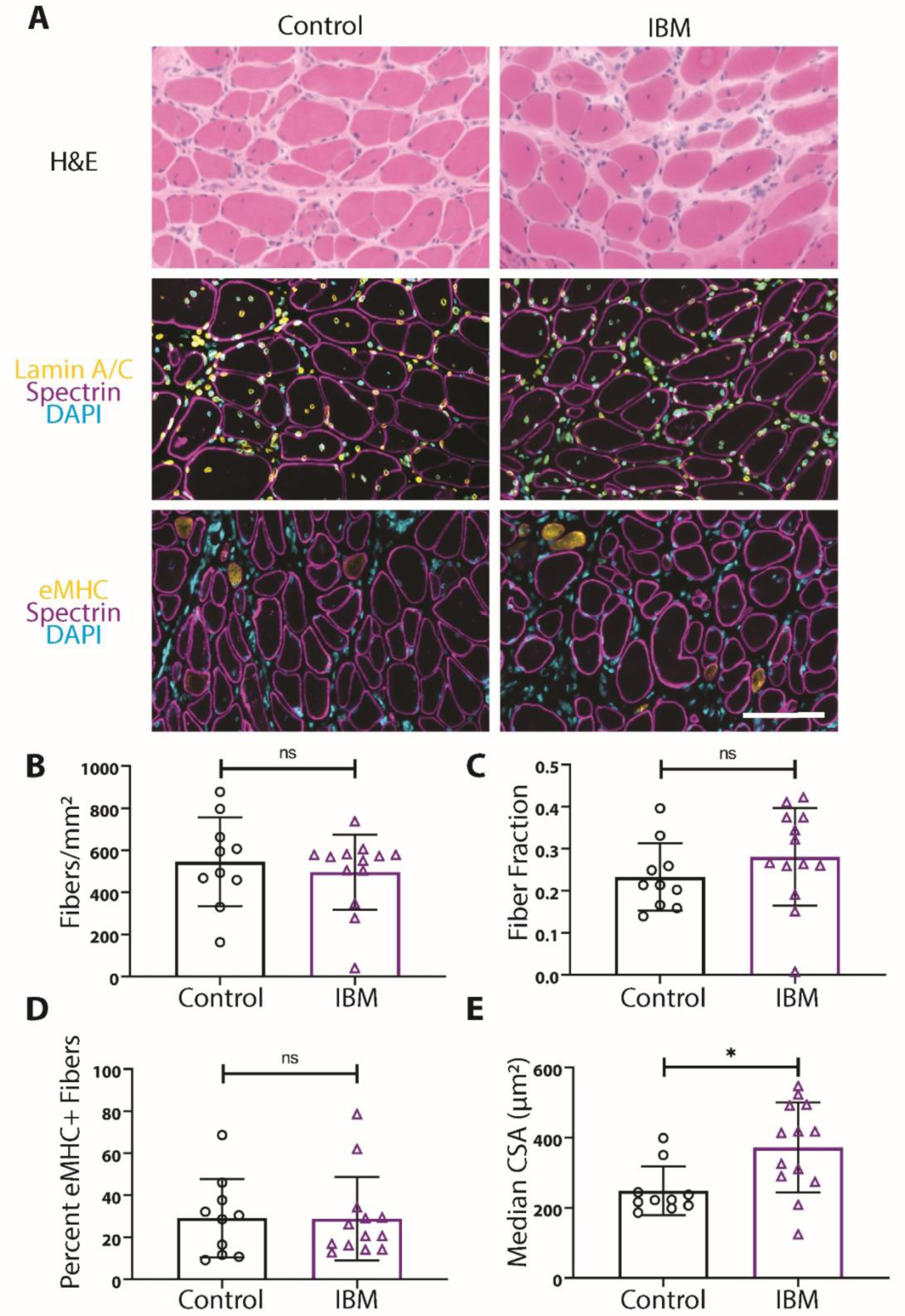
IBM xenografts regenerate robustly in NRG mice. (A) Representative images of 4-month control and IBM xenografts stained with H&E, human spectrin (magenta), human lamin A/C (yellow), embryonic myosin (eMHC) (yellow), and DAPI (cyan). Scale bar shows 100μm. (B-D) The number of fibers per xenograft area (B), the fraction of the xenograft covered by myofibers (C), and the percent of eMHC+ regenerating fibers is similar between control and IBM xenografts (D). (E) The median cross-sectional area (CSA) of myofibers within the xenografts demonstrates a significant increase in fiber size in IBM (*p ≤ 0.05, Mann-Whitney test). For all graphs, each point denotes one patient (control, n = 10; IBM, n = 13).

#### IBM xenografts recapitulate pathological features of the human disease

To determine if newly formed IBM muscle xenografts exhibit features of IBM pathology present in the original biopsy, we first examined 4-month xenografts for rimmed vacuoles, a pathologic hallmark of IBM. Rimmed vacuoles are frequently seen in IBM xenografts as judged by Gomori trichrome (GT) histological stain, whereas they are rarely observed in controls (0.54% vs. 0.05% of fibers; p = 0.007) (**Fig. 3A-B**; **Fig. S5A-C**). Rimmed vacuoles in IBM xenografts show typical histochemical findings, including acid phosphatase positivity and light NADH staining (**Fig. S5B).** Interestingly, the only control xenograft that showed rimmed vacuoles was from a patient diagnosed with a vacuolar myopathy based on muscle biopsy (Case 27) (**Fig. S5A**). These findings demonstrate that regenerated myofibers from IBM patient muscle biopsies develop rimmed vacuoles within a mouse host, indicating that this pathology is likely intrinsic to IBM muscle, rather than resulting from a circulating factor.

**Fig. 3.**
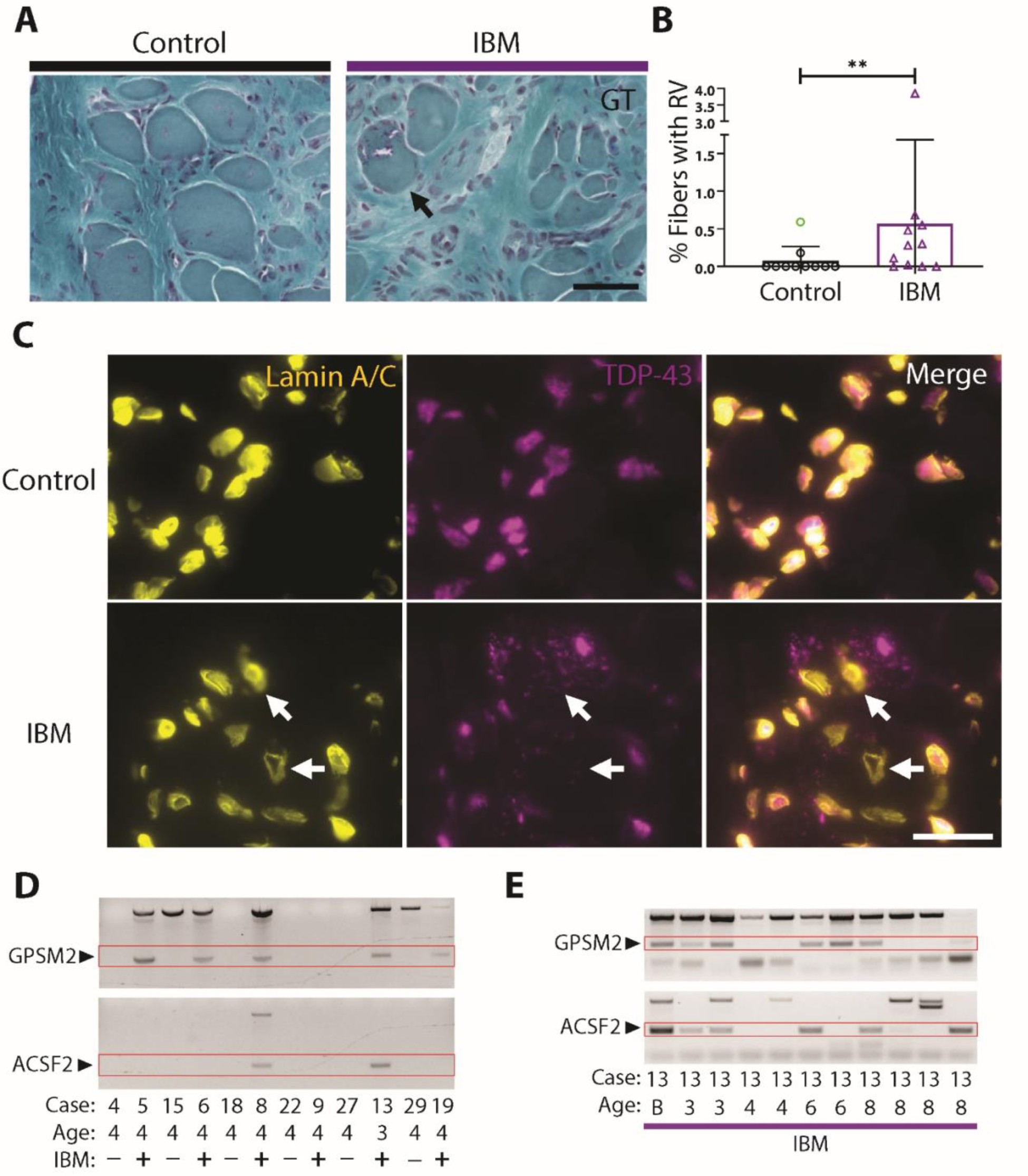
Rimmed Vacuoles and TDP-43 pathology are observed in IBM xenografts. (A) Representative images from Gomori Trichrome stains of 4-month control and IBM xenografts. Examples of rimmed vacuoles (arrow) are exhibited by the IBM xenograft. Scale bar shows 50μm. (B) Quantification of the percent of fibers with rimmed vacuoles (RV) in each group; each point denotes one patient. The control sample highlighted in green indicates the Case 27 patient diagnosed with a vacuolar myopathy (control, n = 10; IBM, n=11). Mann-Whitney test was used to determine significance (*p < 0.05; **p < 0.01). Of note, Case 27 was excluded from the RV statistical analysis as the patient was diagnosed with a vacuolar myopathy. (C) TDP-43 aggregates and examples of nuclear clearing (arrows) can be seen in 4-month IBM xenografts but not in controls. Scale bar shows 25µm. (D) Cryptic exon expression from GPSM2 and ACSF2 was detected in 3- and 4-month IBM xenografts from multiple different cases, but not control xenografts. (E) Cryptic exon expression from TDP-43 target genes GPSM2 and ACSF2 detected in IBM xenografts ranging from 3 to 8-months (B = patient biopsy).

To assess IBM xenografts for pathologic protein aggregation, we first performed immunostaining for p62, an autophagic adaptor protein that binds to ubiquitinated protein aggregates (*20*). Robust p62-positive aggregates are observed at 4-month time points in the majority of IBM xenografts (n = 4/5), but in only one control xenograft (n = 1/5) with a diagnosis of dermatomyositis (**Fig. S6A**). We next looked for evidence of TDP-43 pathology as evidenced by cytoplasmic accumulation and/or nuclear loss of TDP-43. Indeed, nuclear clearance and cytoplasmic aggregation of TDP-43 are observed in 4-month IBM (n = 5/5), but not control (n = 0/5), xenografts (**Fig. 3C**). Having established that the inclusion of TDP-43 cryptic exons occurs in IBM muscle biopsies (**Fig. 1**), we assessed whether TDP-43 splicing repression is compromised in IBM xenografts. We reliably detect cryptic exon expression from TDP-43 target genes GPSM2, ACSF2, and HDGFRP2 within IBM, but not control, xenografts from 2 months to 10 months (**Fig. 3D, E; Table S6**). Cryptic exon incorporation in IBM xenografts corresponds to that of the patient biopsies (i.e., only human biopsies showing the inclusion of cryptic exons result in xenografts with cryptic exon incorporation as shown by IBM cases 9 and 33 (**Fig. 3D**, **Table S3, S6**)). Examination of the inclusion of cryptic exons longitudinally within IBM (cases 8 and 13) reveal that inclusion of cryptic exons can be detected at all time points examined: 3-months (2/2), 4-months (1/4), 6-month (3/4), 8-month (3/7), and 10-month (3/3) (**Fig. 3E**, **Table S6**); approximately 50% (19/37) IBM xenografts show incorporation of cryptic exons regardless of time point, compared with only 3% (1/34) control xenografts. Taken together, these data show that TDP-43 nuclear function is impaired in IBM xenografts similar to what is observed in IBM patient muscle biopsies.

After determining IBM xenografts display characteristic degenerative pathology including rimmed vacuoles and loss-of-TDP-43 function, we next investigated whether the human immune cells transferred via the xenograft surgery persist within regenerated xenografts. Most control xenografts (n = 22/27) recapitulate features of normal muscle tissue including MHC-I staining highlighting capillaries and absence of CD3+ T cells (**Fig. 4A**, **left column**). However, some control xenografts (n = 5/27) and most IBM xenografts (n= 22/33) reveal sarcoplasmic upregulation of MHC-I and the presence of CD3+ T cells (**Fig. 4A, middle and right columns**). Interestingly, multiple human immune cells are present within both control and IBM xenografts, including helper (CD4+) and cytotoxic (CD8+) T cells, B cells (CD20+), macrophages (CD68+), and plasma cells (CD138+) (**Fig. S7**). Despite the presence of CD3+ T cells in both IBM and control xenografts, T cell invasion of non-necrotic fibers is only observed in IBM xenografts (**Fig. 4B**, n = 27 control xenografts and 33 IBM xenografts examined). Importantly, T cells are not observed in regions of mouse muscle adjacent to the human xenograft, arguing against a graft-versus-host response. Furthermore, IBM xenografts show significantly higher numbers of CD3+ T cells than control xenografts (516 vs. 195 per mm^2^, p = 0.01) (**Fig. 4C**), a finding that is also observed in the original muscle biopsies.

**Fig. 4.**
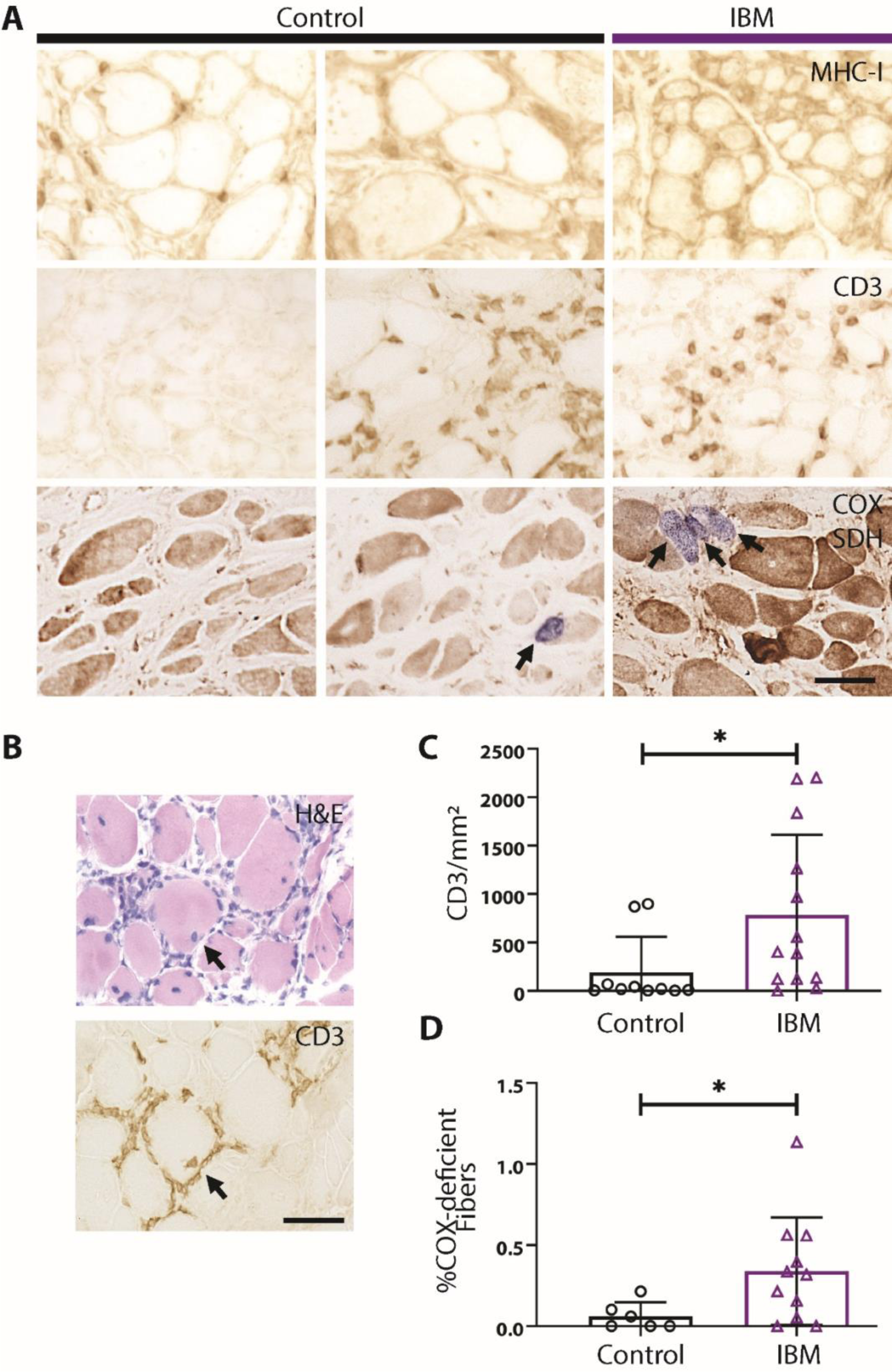
IBM xenografts recapitulate pathological features of human disease. (A) Representative images from MHC-I, CD3, and dual COX/SDH (COX-negative, SDH-dark fibers stain blue) stains of 4-month control and IBM xenografts. As shown, some control xenografts do not have inflammatory cells, whereas others show endomysial immune cells similar to IBM xenografts. (B) Serial sections of a 4-month IBM xenograft stained with H&E and anti-CD3 showing an example of T cell invasion of a non-necrotic fiber. (C) Quantification of the number of CD3+ T cells per xenograft area in 4-month collections. Each point denotes one patient (control, n = 10; IBM, n=13). (D) Quantification of the percent of COX-deficient fibers in each group; each point denotes one patient (control, n = 6; IBM, n=11). Mann-Whitney test was used to determine significance (*p < 0.05; **p < 0.01). All scale bars show 50μm.

Most IBM patient muscle biopsies exhibit mitochondrial pathology, characterized by an increase in the number of COX-deficient fibers (*57–59*), an accumulation of abnormal mitochondria, and mitochondrial DNA (mtDNA) deletions (*60–63*). To assess IBM xenografts for mitochondrial pathology, we carried out dual COX and succinate dehydrogenase (SDH) histological stains on 4-month control and IBM xenografts (**Fig. 4A**). We found that the percentage of COX-deficient fibers was significantly increased in IBM xenografts in comparison to control xenografts (0.34% vs. 0.06% of fibers, p = 0.04) (**Fig. 4D**). Collectively, these results indicate that this xenograft model of IBM successfully recapitulates key features of both degenerative and inflammatory pathology.

#### T cells in IBM xenografts share clones and immunophenotypes with the original IBM biopsy

The majority of IBM patient muscle biopsies show oligoclonal populations of T cells, consistent with the hypothesis that they proliferate in response to unknown antigens (*64–67*). In IBM patient muscle, oligoclonal populations of CD8+ T cells have been shown to persist over time that experience repeated antigen stimulation, as indicated by their loss of CD28 expression, and exhibit a highly differentiated, cytotoxic T cell phenotype, as indicated by a gain of CD57 and KLRG1 expression (*11, 15, 68, 69*). Importantly, we find that IBM xenografts contain both CD57+ and KLRG1+ cells, suggesting that immune cells within xenografts recapitulate immunophenotypes found within the patient muscle (**Fig. 5A**). To better characterize the human immune cells within IBM xenografts, we performed flow cytometry on immune cells isolated from 4-month xenografts (n=4), as well as PBMCs isolated from the same patients at the time of biopsy (n=2) (**Fig. 5B-D**). IBM xenografts show a predominance of CD4+ T cells and large populations of CD57+KLRG1+ CD8+ T cells, similar to what is seen in PBMCs from IBM patients (**Fig. 5C-D**) (*11, 15*). Taken together, these data indicate that the majority of CD8+ T cells in 4-month IBM xenografts contain markers of highly differentiated cytotoxic T cells as seen in IBM biopsies and peripheral blood.

**Fig. 5.**
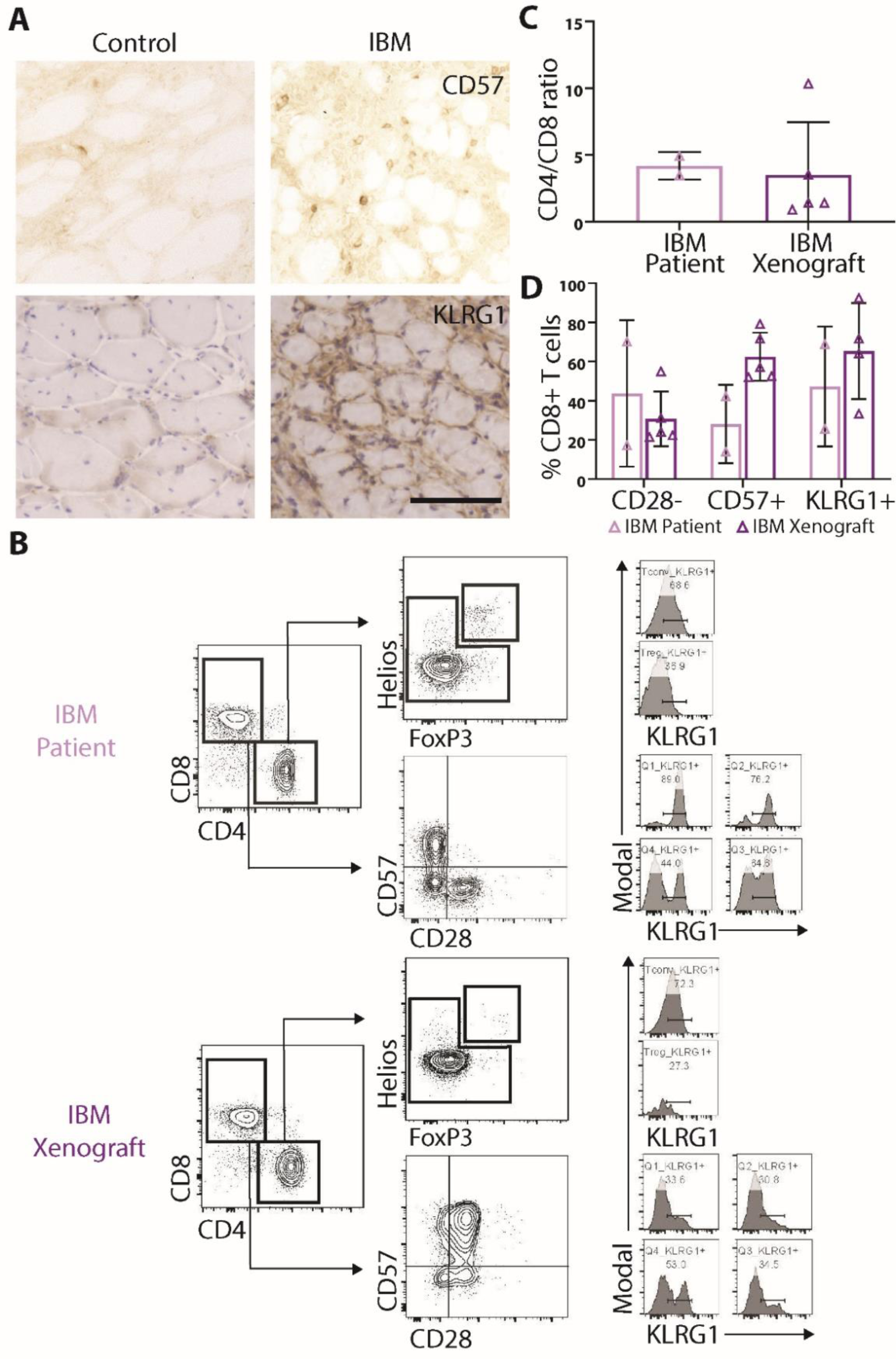
T cells in IBM xenografts are phenotypically similar to IBM patients. (A) Representative CD57 and KLRG1 stains of control and IBM xenografts. Scale bar shows 100µm. (B) Representative Flow Cytometry plots of PBMCs from an IBM patient (top) and a 4-month xenograft (bottom) generated from the same patient. Quantification of (C) CD4/CD8 ratio and (D) percent of CD8+ T cells that are CD28-, CD57+, and KLRG1+ shows similar T cell phenotypes in PBMCs and xenografts.

To explore whether the T cells in IBM xenografts are oligoclonal and share clones in common with the original muscle biopsy, we examined matched biopsy and xenograft T cell receptor (TCR) repertoires using the multiplex PCR-based approach Framework Region 3 AmplifiKation sequencing (FR3AK-seq) (*70*). This analysis reveals that T cells within IBM xenografts are more clonally restricted than the original IBM biopsies (**Fig. 6A**). In addition, IBM xenografts show an increased richness, or number of unique clones, as compared to control xenografts (**Fig. 6B**). Using a Morisita-Horn index to compare the similarity of clones between biopsies and xenografts, we found that T cell clones within a patient biopsy and its corresponding xenografts are more similar to each other than to T cell clones from other IBM patients or xenografts (**Fig. 6C**). Finally, the proportion of T cell clones was compared between IBM patient biopsies and corresponding xenografts at multiple time points (**Fig. 6D**). Multiple xenograft cases showed T cell clones that persist in xenografts across the 1-week to 8-month collections. Together, these data demonstrate that T cells in IBM xenografts are comprised of an enriched subset of clones also detected in the original muscle biopsy.

**Fig 6.**
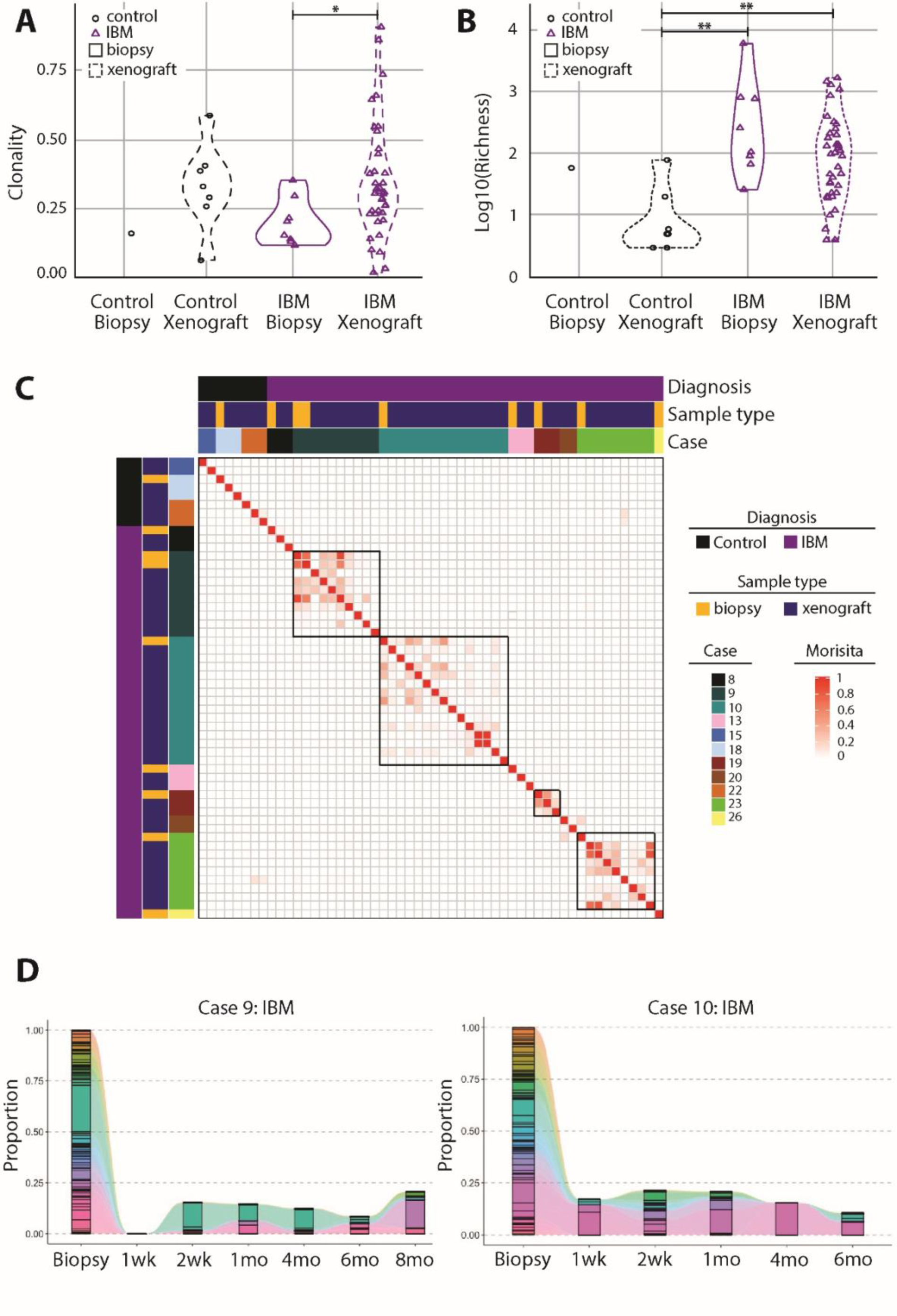
T cells in IBM xenografts show clonality and persistence. T cell receptor sequencing was performed to determine clonality (A) and the number of unique clonotype (richness) (B) in both control and IBM biopsies and xenografts (control biopsy, n=1; control xenograft, n=7; IBM biopsy, n = 8; IBM xenograft, n= 38). This analysis reveals that T cells within IBM xenografts are more clonally restricted than the original IBM biopsies (p = 0.027), and IBM biopsies (p = 0.0031), and xenografts (p = 0.0014) show an increased richness, or number of unique clones as compared to control xenografts. Mann-Whitney test was used to determine significance (*p < 0.05; **p < 0.01). (C) Heatmap displaying the Morisita-Horn index between biopsies and xenografts demonstrates that clones from xenografts of the same IBM biopsy were more similar to each other than to those of xenografts from other IBM biopsies. (D) The proportion of TCR-clones was compared between IBM patient biopsies and corresponding xenografts at multiple time points as shown for two IBM xenograft cases. Each color represents a unique clonotype. These comparisons show that specific clones persist in IBM xenografts over time.

##### Intraperitoneal delivery of OKT3 depletes T cells in IBM xenografts

A central question in IBM is whether inflammation drives pathology or is secondary to muscle degeneration. Recently, highly differentiated effector CD8+KLRG1+ T cells present in IBM muscle have been suggested to be refractory to conventional immunotherapy, and targeted approaches to deplete this subpopulation of T cells are in therapeutic development for IBM (*11, 15*). Monoclonal antibodies can be used to deplete specific immune cells via induction of apoptosis or by antibody-dependent cell-mediated cytotoxicity (ADCC) (*71*). OKT3 was one of the first human monoclonal antibodies created, and it recognizes a nonpolymorphic subunit of the human TCR: CD3ε (*72, 73*). OKT3 has been used clinically to treat transplant rejection and has also been delivered intraperitoneally in xenograft models to ablate human T cells in mice engrafted with human hematopoietic cells (*74*). We hypothesized that treating xenografted mice with OKT3 would specifically eliminate T cells within the grafts and that this would enable us to study the effect of T cells on aspects of IBM pathology.

All four IBM patient biopsies (Cases 23, 26, 36, and 42) used in OKT3 experiments met ENMC clinically-defined IBM criteria and showed marked endomysial inflammation with T cell invasion of non-necrotic fibers and mitochondrial pathology (**Table S4**, **Fig. S8**). Consistent with previous studies showing that about 20% of patients with typical clinical features of IBM do not exhibit rimmed vacuoles (*75, 76*), one of the four IBM cases (Case 36) did not show rimmed vacuoles. Xenografted mice were treated weekly via intraperitoneal injection with 10mg/kg OKT3, as this treatment regimen has been shown to effectively ablate human T cells *in vivo* (*74*). Whereas IBM xenograft mice treated with vehicle alone had an average of 741 CD3+ T cells/mm^2^, OKT3 reduced the number of CD3+ T cells to an average of 93 CD3+ T cells/mm^2^ at 2-months (p = 0.03) and an average of 31 CD3+ T cells/mm^2^ at 4-months (p <0.0001) (**Fig. 7A, B**).

**Fig. 7.**
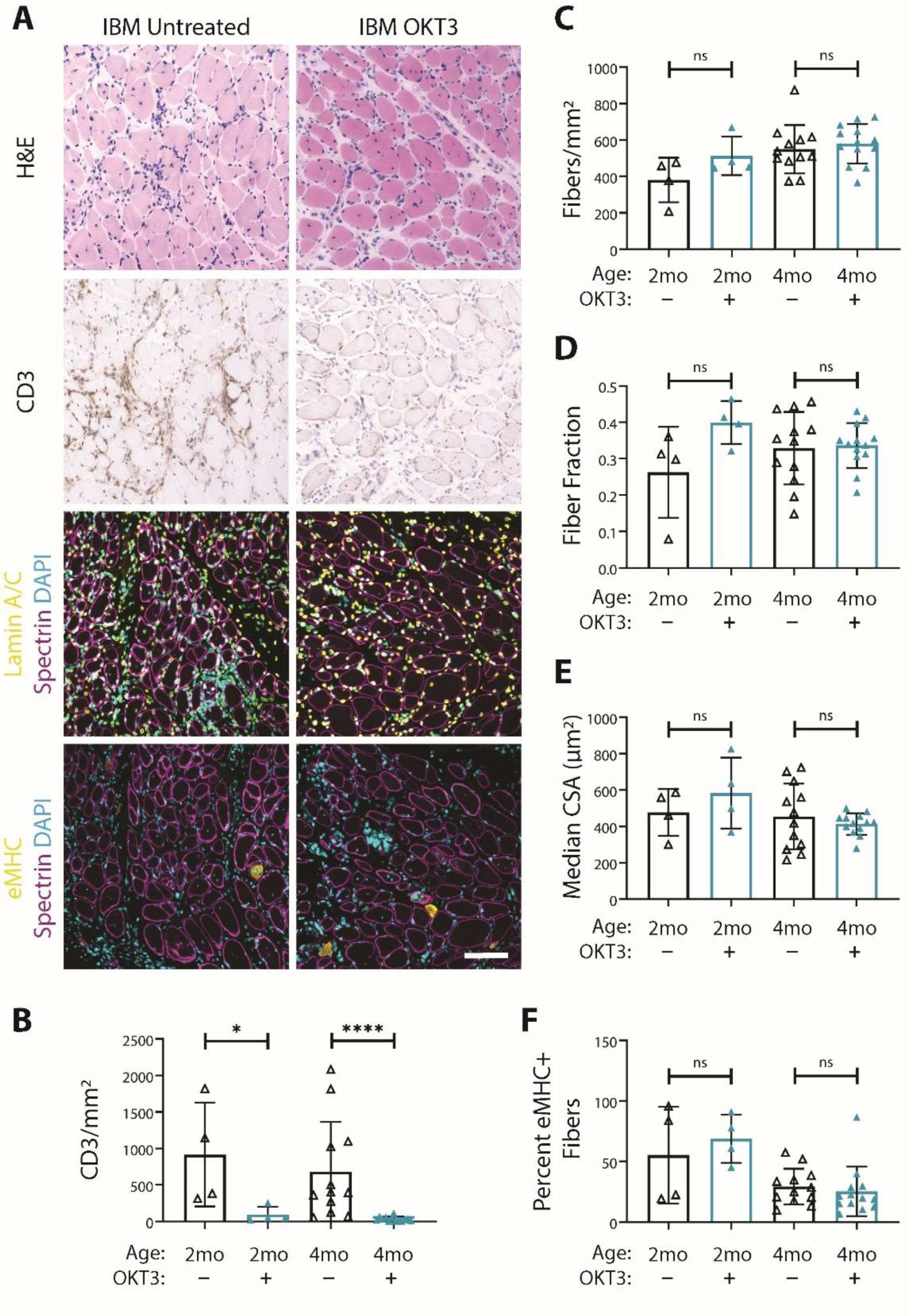
OKT3 treatment ablates T cells from IBM xenografts, but does not impact myofiber regeneration. (A) Representative H&E, CD3, Lamin A/C (yellow), spectrin (magenta), eMHC (yellow), and DAPI stains of 4-month untreated and OKT3 treated IBM xenografts. Scale bar shows 100μm. Quantification of the number of CD3+ T cells over the xenograft area (B) shows the number of T cells is significantly reduced both at 2 months and 4 months by OKT3 treatment. Quantification of the number of fibers over the xenograft area (C), fiber fraction (D), median fiber CSA (E), and percent of eMHC+ fibers (F) show OKT3 treatment did not significantly impact regeneration or fiber morphology. For all graphs, each point denotes one xenograft (2-month untreated n = 4, 2-month OKT3 n = 4, 4-month untreated n = 12, 4-month OKT3 n = 13). Xenografts were obtained from 4 IBM patients (Cases 23,26,36 and 42). Mann-Whitney U test was used to test for significance (*p<0.05, ****p<0.0001).

Although treatment with OKT3 was highly effective in depleting T cells, myofiber regeneration was unchanged between control and treatment groups (**Fig. 7**). The number of regenerated fibers, their median CSA, and the fiber fraction of the xenografts were not significantly affected by OKT3 treatment (**Fig. 7C, D, E**). In addition, OKT3 treatment did not impact the proportion of eMHC+ fibers at 4 months (**Fig. 7F**), indicating that the process of myofiber maturation is unchanged (*56*). In healthy muscle, the inflammatory response to muscle injury is a highly complex and coordinated process involving cells from both the innate and adaptive immune systems (*44*). These data demonstrate that T cells within IBM xenografts do not significantly affect myofiber regeneration in our model.

#### Depletion of T cells in IBM xenografts does not affect rimmed vacuoles or TDP-43 cryptic exons

As xenografts from mice treated with OKT3 or vehicle showed comparable regeneration, these samples are ideal to address the question of how T cells influence aspects of IBM pathology. As expected given then the marked depletion of T cells, OKT3 treatment successfully prevented MHC-I upregulation within xenografts and significantly reduced the number of KLRG1+ cells at 4-months (7.3 vs. 26.3; p = 0.002) (**Fig. 8A, B**). In addition, dual COX/SDH staining showed that OKT3 treatment reduces the number of COX-deficient fibers, although it did not reach statistical significance (from 0.37 to 0.19%, p = 0.12) (**Fig. 8A, C**). These data suggest that T cells are required for mitochondrial pathology in IBM xenografts. Indeed, we find that the number of COX-deficient fibers is significantly correlated to the number of CD3+ T cells in IBM xenografts (Control R^2^: 0.03; IBM R^2^: 0.2; p = 0.02) (**Fig. 8D**), consistent with previous work demonstrating positive correlations between the number of COX-deficient muscle fibers and the severity of inflammation in IBM patient biopsies (*77*). Interestingly, this relationship to IBM xenografts is not observed in the few control xenograft samples that showed similar numbers of CD3+ T cells (**Fig. 8D**). Taken together, the amelioration of MHC-I upregulation and reduction in mitochondrial pathology achieved by OKT3 treatment suggest that T cells are required for both of these pathological features in IBM.

**Fig. 8.**
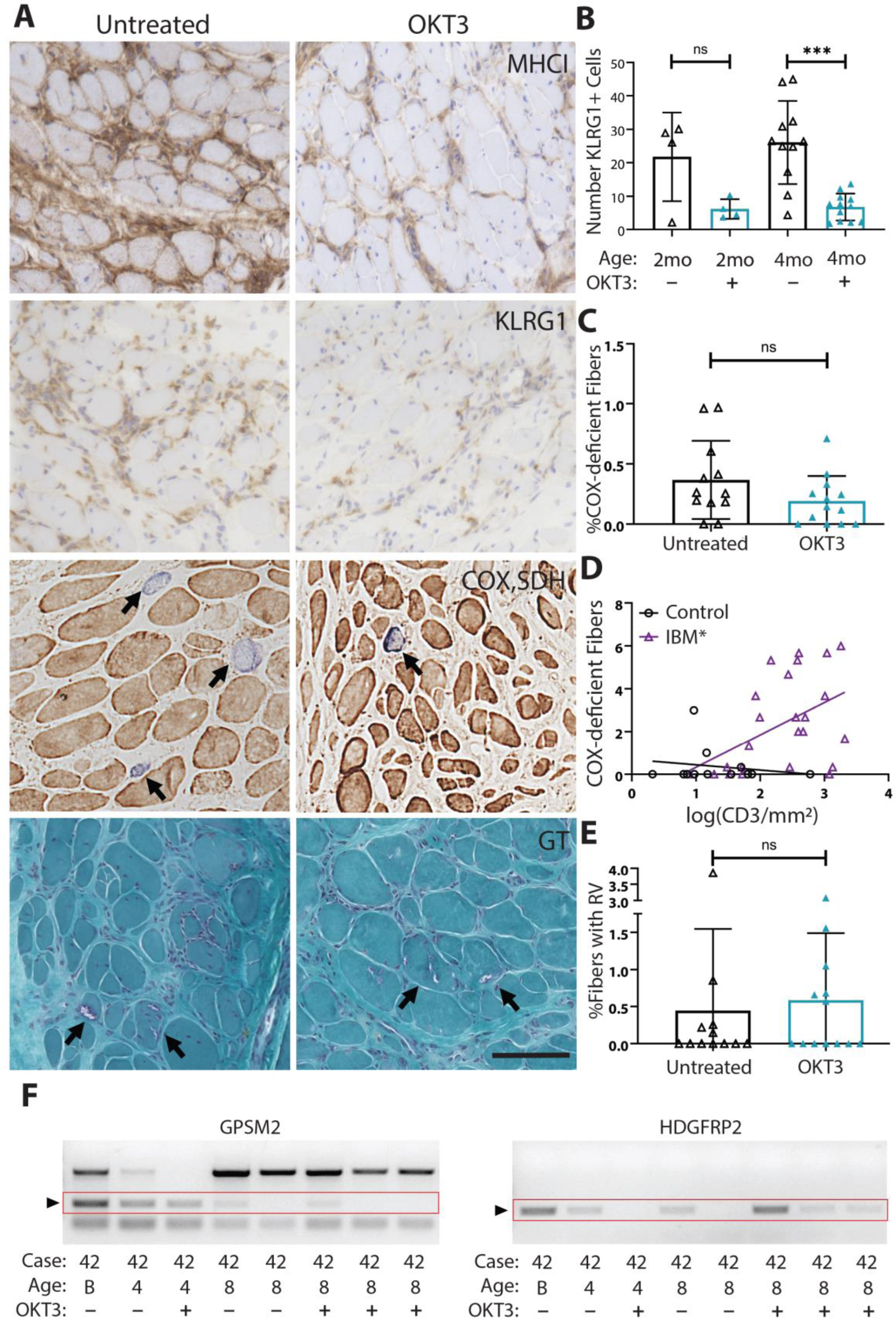
OKT3 depletion of T cells reduces mitochondrial pathology and inflammation but not rimmed vacuoles or TDP-43 pathology. (A) Representative images from MHC-I, KLRG1, dual COX-SDH, and GT stains of untreated and OKT3 treated 4-month IBM xenografts. COX-deficient fibers appear light to dark blue (arrows) and both untreated and OKT3 treated xenografts have rimmed vacuoles (arrows). Scale bar shows 100µm. (B) Quantification of the number of KLRG1+ cells in untreated and OKT3 treated xenografts at both 2- (untreated n = 4, OKT3 n = 4) and 4-month (untreated n = 11, OKT3 n =12) time points. Mann-Whitney test was used to determine significance (***p<0.001). (C) Quantification of the percent of COX-deficient fibers in 4-month xenografts, each point denotes one xenograft (untreated n = 12, OKT3 n =13) and Mann-Whitney test was used to determine significance (p = 0.16). (D) Correlation of the number of CD3+ T cells to the number of COX-deficient fibers in 4-month xenografts, each point denotes one xenograft (control, n = 12; IBM n=24). Control xenografts were obtained from 7 patients and IBM xenografts were obtained from 11 patients. The Pearson correlation coefficient was used to determine the degree of linear correlation in each group (Control, r = −0.18, p = 0.57; IBM, r = 0.47, *p = 0.02). (E) Quantification of the percent of myofibers containing rimmed vacuoles (RV) in untreated and OKT3 treated xenografts at 4 months (untreated n = 12, OKT3 n =13). Mann-Whitney test was used to determine significance. (F) Cryptic exon expression from TDP-43 target genes GPSM2 and HDGFRP2 can be detected in both untreated and OKT3 treated IBM xenografts.

Given that T cell depletion significantly reduced the inflammatory changes in IBM xenografts, we next determined whether OKT3 affected degenerative muscle pathology. As determined by Gomori Trichrome staining, OKT3 treatment did not affect the percent of myofibers containing rimmed vacuoles (**Fig. 8A, E)**. Similarly, GPSM2, ACSF2, and/or HDGFRP2 cryptic exons were detected in OKT3-treated IBM xenografts at 2-, 4- and 8-month time points (**Fig. 8F**, **Table S6**). These data suggest that rimmed vacuole pathology and loss of nuclear TDP-43 function within myofibers do not require T cells. Furthermore, by comparing IBM xenografts with and without cryptic exon expression, we find no difference in CD3+ T cell number (**Fig. S9E**), but we do find that xenografts with cryptic exon expression have a significantly higher percentage of fibers with rimmed vacuoles (1.2 vs 0.2%, p < 0.008), suggesting that rimmed vacuoles are associated with loss of TDP-43 function (**Fig. S9**). These findings demonstrate that rimmed vacuole pathology and TDP-43 dysfunction in IBM xenografts persist in IBM xenografts depleted of T cells.

### DISCUSSION

The intertwined degenerative and inflammatory pathological features have fueled the debate underlying the pathogenesis of IBM and have impeded the generation of laboratory models. While IBM muscle biopsies exhibit nuclear clearance and cytoplasmic aggregation of TDP-43, it is not clear whether loss of nuclear TDP-43 function occurs in this disease. Evidence that loss of TDP-43 function is a common cellular pathogenic mechanism in neurodegenerative disease exhibiting TDP-43 pathology would support such a view. Indeed, recent evidence strongly supports the view that loss of TDP-43 splicing repression triggers neurodegeneration: 1) TDP-43 splicing repression is compromised in brains of ALS-FTD and Alzheimer’s Disease (*26*); 2) TDP-43 nuclear depletion in brain neurons has been reported at the pre-symptomatic stage in a *C9orf72* linked FTD-ALS patient (*78*), suggesting that loss of splicing repression represents an early event in disease progression; 3) ALS-linked mutant TDP-43 fails to repress non-conserved cryptic exons independent of TDP-43 cytoplasmic aggregation (*28, 29*), and facilitates the formation of RNA-free TDP-43 into anisotropic intranuclear liquid spherical shells (*79*). Our demonstration of TDP-43 cryptic exons in IBM patient muscle (**Fig. 1**) is consistent with the notion that nuclear depletion of TDP-43 represents an early contributor to IBM pathogenesis. While many different immunohistochemical assays and combinations of clinical and pathological features have been suggested to have high sensitivity and specificity for the diagnosis of IBM, the PCR-based cryptic exon detection assay we report here is the first functional assay to demonstrate high sensitivity (84%) and specificity (99%) for IBM diagnosis in a large myositis cohort (119 patients: IBM, n =44; Control, n=75). Since the incorporation of some cryptic exons (those that are fused in-frame) into mRNAs to encode novel epitopes (neoantigens), we hypothesize that such neoantigens may contribute to the autoimmune response in IBM. Furthermore, detection of these neoantigens in serum or muscle could be used as functional biomarkers for clinical applications.

Here, we show skeletal muscle xenografts provide a new approach to model acquired muscle diseases such as IBM. A variety of laboratory models have been developed for various forms of hereditary inclusion body myopathy (hIBM), such as transgenic (*80, 81*) or knock-in models (*82*) of IBMPFD caused by mutations in VCP. However, while hIBMs share degenerative pathological features (TDP-43 pathology, protein aggregates, and rimmed vacuoles) with IBM, they are clinically distinct from IBM in that patients have an earlier age of onset and different patterns of muscle involvement, and in stark contrast with IBM, muscle biopsies typically lack inflammation.

In addition, other animal models have been developed to recapitulate specific features of IBM (*83*). For example, transgenic approaches that drive β-amyloid (Aβ) expression in skeletal muscle results in a vacuolar myopathy (*84, 85*). However, these models also fail to show inflammatory pathology, and the potential contribution of β-amyloid and amyloid precursor protein (APP) in the pathogenesis of IBM remains controversial (*86, 87*). While transgenic mice that conditionally overexpress MHC-I show myofiber degeneration, they lack other aspects of IBM pathology (*88*). Thus, existing mouse models of IBM recapitulate some aspects of IBM pathology, but none show the full spectrum of pathological features. The xenograft model described here is the first animal model of IBM to recapitulate both degenerative and inflammatory hallmarks of the disease. For therapeutic development in rare or sporadic diseases that often lack laboratory models, xenograft models of disease have increasingly proven to be valuable tools. These xenografts can recapitulate the complex genetic and epigenetic abnormalities that exist in human disease that may never be reproducible in other animal models, and xenografts form a complete *in vivo* system for modeling disease.

Our data show that IBM patient muscle robustly regenerates in immunodeficient mice to form skeletal muscle xenografts despite the presence of an inflammatory milieu. Importantly, the characteristic degenerative pathological features of IBM are recapitulated in this xenograft model. First, most IBM xenografts display robust rimmed vacuoles at 4-months, and this pathological feature was only observed in xenografts from one control case in a patient with a suspected genetic vacuolar myopathy. Second, nuclear clearance of TDP-43 leading to missplicing and incorporation of cryptic exons is detected in the majority of IBM xenograft cases, indicating a loss of TDP-43 function in our model. Since all mature myofibers cut during the biopsy procedure are replaced with myofibers that form *de novo* from satellite cells by 4-months, our data indicate that rimmed vacuole formation and TDP-43 pathology are intrinsic to IBM patient muscle and do not require any factors circulating within patient blood. TDP-43 functions in normal muscle regeneration through the formation of cytoplasmic, amyloid-like “myo-granules,” which associate with sarcomeric mRNAs and localize to sites of sarcomere formation (*89*). These myo-granules form in healthy muscle following injury and are readily cleared as myofibers mature. However, purified myo-granules can seed the formation of amyloid-like fibrils *in vitro*, and therefore may lead to the formation of stable aggregates that may drive disease pathology (*90*), an idea that can now be tested in our IBM xenograft model.

In addition to these degenerative features, IBM xenografts also show elevation of MHC-I, intense endomysial inflammation, and oligoclonal expansion of CD8+ T cells that express markers of highly-differentiated cytotoxic T cells including CD57 and KLRG1. The persistence of these T cells and evidence of invasion of non-necrotic myofibers in IBM xenografts strongly suggests ongoing antigen stimulation by newly forming myofibers. In contrast, there is no evidence of graft-versus-host disease, as human T cells do not invade surrounding mouse muscle, and the expanded T cell clones in xenografts mirror those of the original IBM muscle biopsy.

Overall, this xenograft model of IBM is the first comprehensive laboratory model that exhibits both degenerative and inflammatory features. However, this model does have several limitations. First, typical behavioral studies used to assess muscle weakness, such as the rotarod test or the Morris water maze will be challenging to apply to this model (*91*). However, xenograft contractility can be assessed *in vitro* and individual myofibers can be isolated to study calcium release dynamics (*37*). In our studies, the number of xenografts that could be obtained from each individual patient was limited by the size of the additional research piece obtained at diagnostic biopsy; however, many more xenografts could be performed from a single patient if a dedicated research muscle biopsy was performed on a relatively preserved muscle. Given the variability of both inflammatory and degenerative features observed in IBM xenografts (similar to that seen in different muscle biopsies from the same IBM patient), research studies using this model to test therapies will benefit from increasing the number of xenografts performed from a single patient– while this may not be practical in patients with advanced-stage disease, this can readily be performed in patients with early disease and relatively preserved muscle bulk.

Using a monoclonal CD3 antibody (OKT3) (*73, 74, 92*), we successfully depleted 96% of T cells from IBM xenografts. This treatment normalized MHC-I expression in myofibers, demonstrating that T cells are required for MHC-I upregulation. That T cell burden correlates with the number of COX-deficient fibers in IBM xenografts, as has been identified in IBM biopsies, and that OKT3-mediated T cell ablation suppresses these mitochondrial defects suggest that T cells directly contribute to mitochondrial pathology in IBM. Such a view is consistent with *in vitro* studies showing primary myotubes cultured with pro-inflammatory cytokines such as interferon-gamma (IFN-γ) and interleukin-1beta (IL-1β) led to a decrease in levels of COX protein (*93*).

While OKT3 treatment significantly ameliorated inflammatory changes in IBM xenografts, degenerative pathological features including rimmed vacuoles and loss of TDP-43 function persist. This finding may explain why IBM patients do not respond to immunosuppressive treatment. Although we cannot exclude the possibility that T cells induce these degenerative features during early myofiber regeneration, prior to OKT3-mediated ablation, our data are most consistent with a model in which loss of TDP-43 function and rimmed vacuole formation in IBM occur independently of T cells. These findings support the view that IBM should be considered within the spectrum of TDP-43 proteinopathy, along with ALS, FTD, and other neurodegenerative diseases exhibiting TDP-43 pathology, and that therapeutic development should focus on both correction of proteostasis defects and restoring TDP-43 function as well as ameliorating endomysial inflammation.

### MATERIALS AND METHODS

#### Human Muscle Biopsy

All use of research specimens from human subjects was approved by the Johns Hopkins Institutional Review Board (IRB) to protect the rights and welfare of the participants. Patients with suspected muscle disease scheduled for a diagnostic muscle biopsy provided informed consent prior to providing an extra muscle sample for use in xenograft surgeries. More specifically, for inclusion in this study, IBM cases met ENMC 2011 criteria for clinically-defined or probable IBM (*4*). Control samples were selected from non-IBM patients, including dermatomyositis, polymyositis, immune-mediated necrotizing myopathy, as well as patients with normal biopsies or non-inflammatory pathological features. Patient samples with excessive fibroadipose replacement or in poor condition were excluded. Under sterile conditions in the operating room, approximately one gram of tissue was removed from muscles having a strength of ≥4 (MRC scale (*94*). This tissue was then dissected into approximately 7 x 3 x 3mm strips of longitudinal fibers and taken immediately to the animal suite for xenografting.

#### Animal Husbandry

All animal experiments were approved by the Johns Hopkins University Institutional Animal Care and Use Committee (IACUC) in accordance with the National Institutes of Health (NIH) Guide for the Care and Use of Laboratory Animals. Male NOD-Rag1[null] IL2rγ[null] (NRG) mice (The Jackson Labs, stock 007799) were used for all experiments, and mice were provided water and an irradiated antibiotic diet (Envigo, TD.06596) ad libitum. For xenografted mice treated with 10mg/kg OKT3 (Fisher, 50561956), stock OKT3 was diluted with sterile PBS and injected intraperitoneally immediately after the xenograft surgery and once weekly until xenograft collection was performed. Control “untreated” mice in OKT3 experiments were injected with sterile PBS following the same treatment regimen.

#### Xenograft Procedures

This xenograft procedures were performed as previously described (*55*). Briefly, 8- to 12-week NRG mice were anesthetized with 1.5% isoflurane, and 0.1mg/kg Buprenorphine (ZooPharm) was administered subcutaneously for pre-emptive analgesia. The tibialis anterior and extensor digitorum longus muscles were removed, and a 7 x 3 x 3mm strip of dissected human muscle was placed in the tibial compartment and ligated with non-absorbable sutures (6-0 Surgipro, Covidien) to the tendons of the peroneus longus muscle. The incision was closed with surgical glue (Histoacryl, Tissue seal) and stainless-steel wound clips (AutoClip System, F.S.T). The analgesic Carprofen (Rimadyl, Patterson Veterinary) was given subcutaneously after the surgery at a dose of 5mg/kg. To harvest the muscle, NRG host mice were anesthetized with 1.5% isofluorane, the leg was shaved, and the skin overlying the xenograft was opened and the location of the graft determined by the position of the non-absorbable sutures. The xenograft was then removed, snap-frozen in 2-methylbutane (Fisher, 03551), and sectioned at 10 µm using a cryostat (Leica, CM1860 UV). Following collection, the NRG host mice were euthanized consistent with AVMA guidelines.

#### Histochemical Staining

Fresh, 10µm sections were rehydrated through an ethanol dilution series: three 3-minute washes in 100% ethanol (Fisher, BP2818100), a 3-minute wash in 95% ethanol, and a 3-minute wash in 80% ethanol). Following a 5-minute wash in distilled water (dH2O), slides were placed in Hematoxylin (Poly Scientific, s212A) for 3 minutes, washed quickly in dH2O, and placed in Tap water for 5 minutes to allow the stain to develop. Slides were dipped 12 times in acid ethanol (1 mL concentrated Hydrochloric Acid (Fisher, SA48) in 400mL 70% ethanol), rinsed twice in Tap water for 1 minute, and placed in distilled water for 5 minutes. Slides were then placed in Eosin (Poly Scientific, s176) for 30 seconds, and dehydrated in ethanol. Slides were cleared in xylene (Fisher, X5) and mounted using Permount (Fisher, SP15). Gomori-Trichrome, NADH, Acid Phosphatase staining, and dual COX/SDH was performed as previously described (*95*).

#### Immunohistochemistry

10µm fresh frozen sections were fixed with ice-cold methanol (Fisher, A412) for 10 minutes and blocked with anti-mouse IgG (MKB-2215, Vector Laboratories) or with a blocking solution consisting of 2% normal goat serum in PBS. The primary antibodies used were: anti-human spectrin (NCL-SPEC1, Leica, 1:50), anti-human lamin A/C (Abcam, Ab40567, 1:50), anti-human MHC-1 (SC-32235, Santa Cruz, 1:300), anti-human p62 (SC-25575, Santa Cruz, 1:250), anti-SQSTM1/p62 (D-3) (sc-28359,1:200), anti-TDP-43 (Proteintech 10782-2-AP and 12892-1-AP, 1:1000), anti-embryonic Myosin Heavy Chain (eMHC, MYH3) (F1.652, DSHB, 1:10), anti-CD3 mouse (M725401-2, DAKO, 1:60), anti-CD3 rabbit (A0452, DAKO, 1:60), anti-human CD4 (ab133616, abcam, 1:100), anti-CD8 (M710301-2,DAKO, 1:60), anti-Ki-67 (ab92742, abcam,1:60), anti-CD20 (M0755, DAKO, 1:200), anti-CD68 (M0718, DAKO, 1:60), anti-CD138 (M7228, DAKO, 1:100). The secondary antibodies used were AlexaFluor 488 goat anti-mouse IgG1, AlexaFluor 594 goat anti-mouse IgG2b, and AlexaFluor 594 goat anti-mouse IgG1a (all Life technologies, 1:500). Biotinylated Goat Anti-Rabbit IgG Antibody (BA-1000, Vector Laboratories, 1:100) and Biotinylated Goat Anti-Mouse IgG Antibody (BA-9200, Vector Laboratories, 1:100) were used for DAB peroxide staining (SK-4100, Vector Laboratories). Where applicable, nuclei were labeled with DAPI in mounting medium (P36931, Invitrogen).

#### Cell culture and manipulation

Human skeletal myoblast cells were obtained from Zen-Bio (SKB-F) and cultured in Skeletal Muscle Cell Growth Medium (Zen-Bio, SKM-M). Knockdown of TDP-43 was performed by transfecting siRNA targeting the TARDBP transcript (Sigma-Aldrich, EHU109221) while control was transfected with negative control siRNA (Life Tech., 4390843). Transfection of siRNA was achieved using Viromer Blue (Lipocalyx, VB-01LB-01).

#### TDP-43 Cryptic Exon Detection

RNA was extracted from xenograft samples and human biopsies using TRIzol (Fisher Scientific, 15596018). A cDNA library was prepared using Protoscript II First Strand cDNA synthesis kit with random primers (NEB, E6560L), and cryptic exons were PCR amplified using the Dreamtaq Kit (Fisher, K1081P) following the protocol described below. Primer sequences for TDP-43 target genes are summarized below. PCR products were visualized via gel electrophoresis on a 2% agarose (Fisher, 17850) gel containing 0.5µg/mL Ethidium bromide (Fisher, A25645).

Protocol for Cryptic Exon Product Amplification

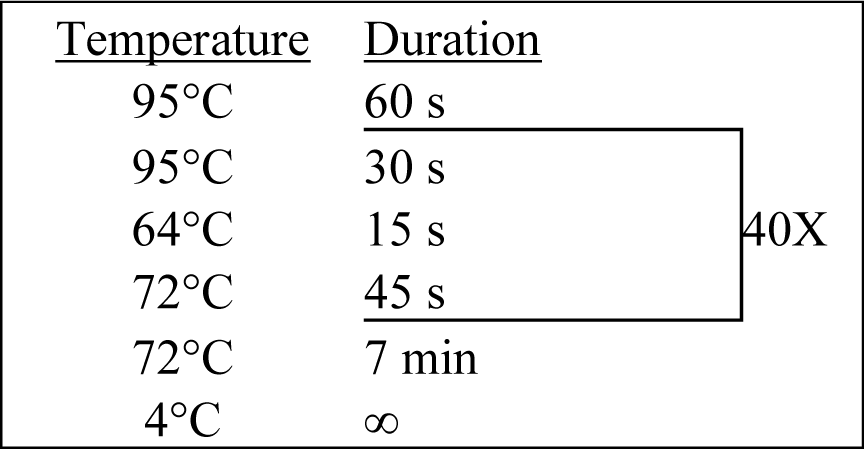

Primer Sequences for Cryptic Exon Detection

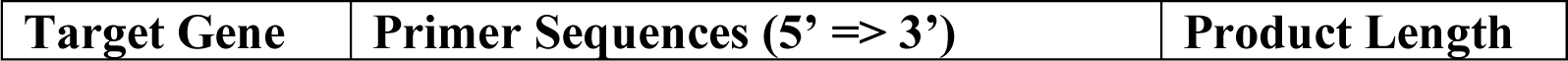
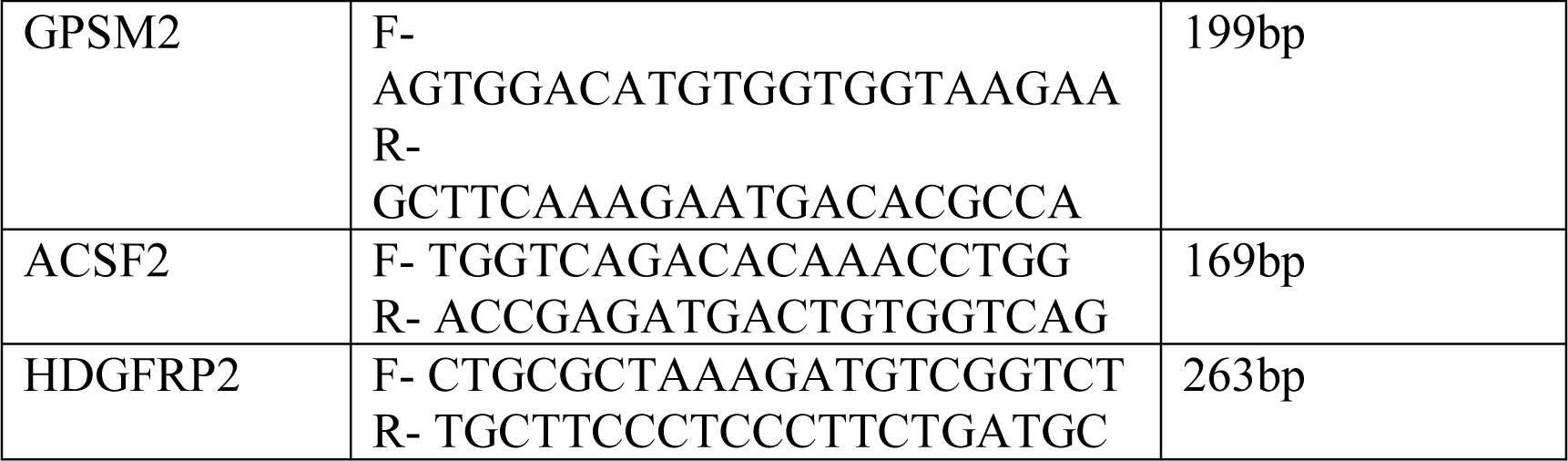

#### T cell Receptor Sequencing

T cell Receptor (TCR) sequencing was performed using Framework Region 3 (FR3) AmplifiKation sequencing (“FR3AK-seq”) as previously described (*70*). Briefly, RNA was TRIzol extracted from human biopsies and xenograft samples, and 1000 cellular equivalents of monoclonal Jurkat RNA was added to each sample as a spike-in. This RNA was then reverse-transcribed using a TCR beta (TCRB) chain constant region reverse primer with Superscript III First-Strand Synthesis System (Invitrogen). Forward primers targeting TCRB FR3 and a nested reverse TCRB constant region primer were then used to PCR amplify the TCRB cDNA with KAPA2G Fast Multiplex Mix (Roche). A second PCR was then performed to add appropriate Illumina sequencing adapters using Herculase II Fusion DNA polymerase (Agilent). Libraries were quantified using the KAPA Library Quantification Kit for Illumina Platforms (Roche). Sequencing was performed on a HiSeq 2500, and results were analyzed using MiXCR v2.1.11 (*96*). Clonality was calculated as (1 – Shannon’s equitability) (*97*). Immunarch (*98*) was used to calculate richness, Morisita’s overlap index, and to quantify public clones. R software (www.r-project.org) was used for statistical analyses and plot generation.

#### Flow Cytometry

Flow Cytometry experiments were performed as previously described (*99*). To eliminate non-muscle residing immune cells, mice were perfused with PBS and lymph nodes were removed. Single-cell suspensions were prepared from the isolated xenografts and incubated with anti-CD16/32 (clone 2.4G2) to block Fc receptors. These suspensions were then stained with human antibodies for CD3, CD4, CD8, CD28, CD57, and DAPI or Blue-Fluorescent Reactive Dye (Invitrogen Life Technologies) to assess cell viability. The analysis was performed on live cells on a BD LSRII flow cytometer with FACSDiva software (BD Bioscience). Post-acquisition analysis was performed with Flowjo software version 9.1.

#### Microscopy and Image Analysis

Fluorescent and transmitted light microscopy was carried out at the Johns Hopkins NINDS Multiphoton Imaging Core on a Keyence (BZ-X700) widefield, inverted microscope. Image analysis was performed in Fiji (*100*). Analysis of fiber cross-sectional area was semi-automated using MuscleJ (*101*).

#### Statistical analysis

All statistical analyses were performed using GraphPad Prism version 8.3.1 (GraphPad Software, La Jolla California USA, www.graphpad.com). For non-normally-distributed data (Shapiro-Wilk test, p < 0.05), the nonparametric Mann-Whitney test was used to determine significance between two groups, or Fisher’s exact test where noted. Data are presented as means +/− SD unless otherwise indicated in figure legends. Significance markers on figures are from *post hoc* analysis (ns, not significant; *p ≤ 0.05, **p ≤ 0.01; ***p ≤ 0.001; ****p ≤ 0.00001) with values of *p* ≤ 0.05 considered significant unless otherwise noted in figure legends.

## Supplementary Materials

Fig. S1: Consensus sequences flank cryptic exons targeted by TDP-43

Fig. S2: Sequence and alignment validation of cryptic exon RT-PCR: ACSF2 and GPSM2

Fig. S3: Sequence and alignment validation of cryptic exon RT-PCR: HDGFRP2 and ZFP91

Fig. S4: Description of Xenograft Model

Fig. S5: Rimmed vacuoles in IBM xenografts

Fig. S6: IBM xenografts show p62 aggregation at 4 months

Fig. S7: Multiple human immune cells are present within xenografts

Fig. S8: Histological features of human biopsies in OKT3 experiments

Fig. S9: Rimmed vacuoles are more frequent in xenografts containing cryptic exons.

Table S1: TDP-43 associated cryptic exons in human myoblasts.

Table S2: Demographics, Clinical, and muscle biopsy findings in Cryptic Exon Patient Cohort

Table S3: Patient cohort cryptic exon expression

Table S4: Clinical, demographic, and biopsy details of patients in the Xenograft Cohort

Table S5: Summary of the sex ratio, age, disease duration, and biopsy location of the Xenograft Patient Cohort.

Table S6: Xenograft cryptic exon expression summary

## Acknowledgments

We thank the staff of the Next Generation Sequencing Center (JHMI) for their RNA sequencing service, and the Johns Hopkins NINDS Multiphoton Imaging Core (NS050274) for their technical expertise and providing imaging equipment.

## Author contributions

K.A.B., J.P.L., K.E.B., P.C.W., and T.E.L. designed experiments and interpreted results. K.A.B., J.P.L., K.E.B., K.A.R., A.W., C.I., W.T., and N.R. carried out experiments. J.M.M and H.B.L performed FR3AK-sequencing experiments and analysis. K.R.W provided technical expertise and training in the xenograft surgery. L.W.O. and A.M.C. performed human biopsies. J.M.K. and S.A.V. performed flow cytometry experiments and analysis. I.P-F. and A.L.M. provided key intellectual contributions. K.A.B., P.C.W. and T.E.L. wrote the manuscript, which was approved by all authors.

