## Supplemental figures and tables for "Loss of TDP-43 function and rimmed vacuoles persist after T cell depletion in a xenograft model of sporadic inclusion body myositis"

Fig. S1: Consensus sequences flank cryptic exons targeted by TDP-43

Fig. S2: Sequence and alignment validation of cryptic exon RT-PCR: ACSF2 and GPSM2

Fig. S3: Sequence and alignment validation of cryptic exon RT-PCR: HDGFRP2 and ZFP91

Fig. S4: Description of Xenograft Model

Fig. S5: Rimmed vacuoles in IBM xenografts

Fig. S6: IBM xenografts show p62-positive aggregates

Fig. S7: Multiple human immune cells are present within xenografts.

Fig. S8: Histological features of human biopsies in OKT3 experiments

Fig. S9: Rimmed vacuoles are more frequent in xenografts containing cryptic exons.

Table S1: TDP-43 associated cryptic exons in human myoblasts.

Table S2: Demographics, Clinical, and muscle biopsy findings in Cryptic Exon Patient Cohort

Table S3: Patient cohort cryptic exon expression

Table S4: Clinical, demographic, and biopsy details of patients in the Xenograft Cohort

Table S5: Summary of the sex ratio, age, disease duration, and biopsy location of the Xenograft Patient Cohort.

Table S6: Xenograft cryptic exon expression summary

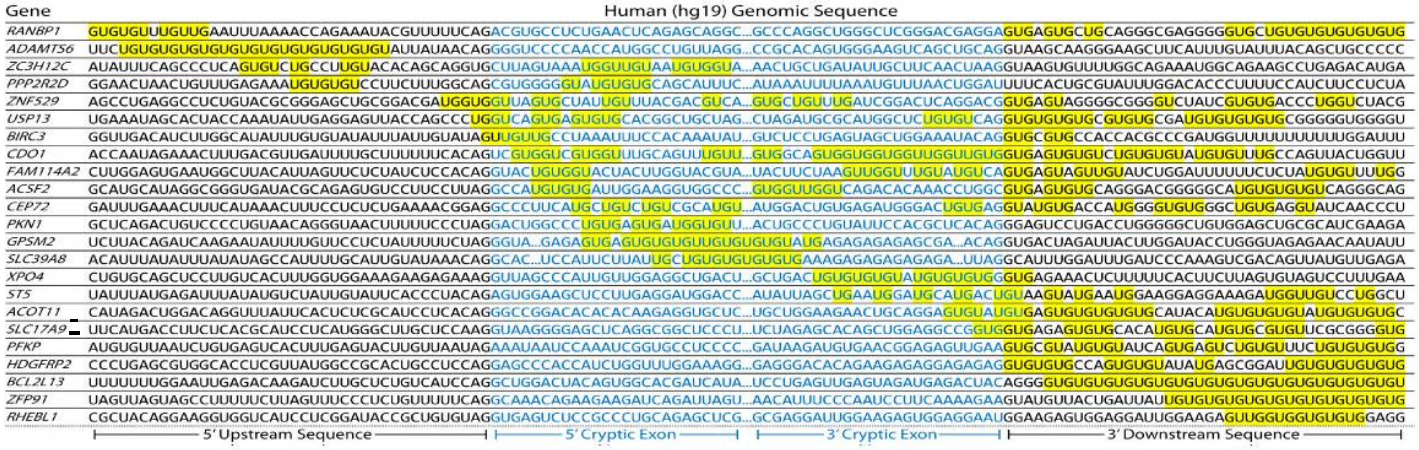

**Fig. S1: Consensus sequences flank cryptic exons targeted by TDP-43**. Cryptic exons are flanked by UG tandem repeats that exist upstream, downstream, or internally in TDP43 KD myoblast cells.

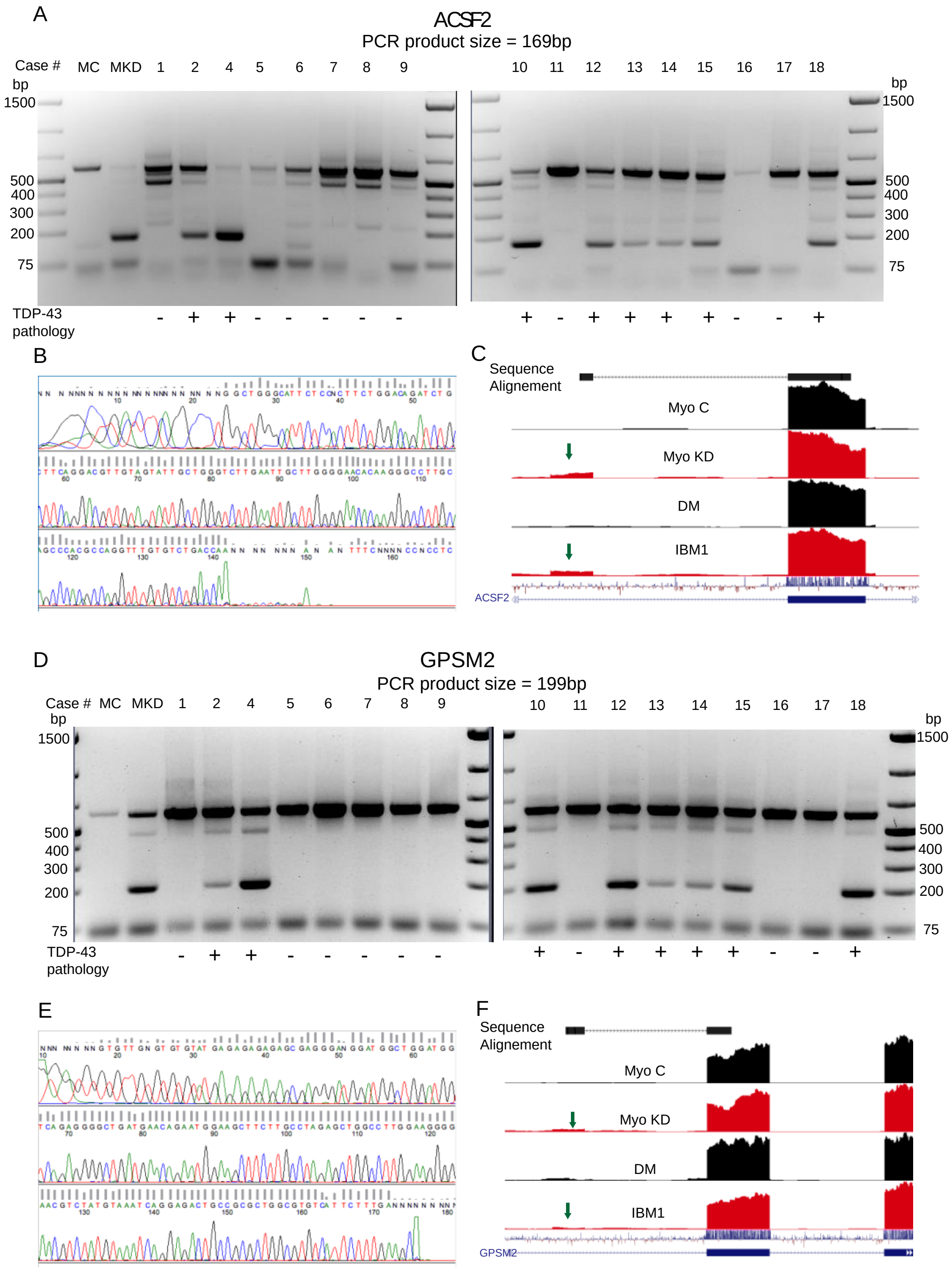

**Fig. S2. Sequence and alignment validation of cryptic exon RT-PCR: ACSF2 and GPSM2**

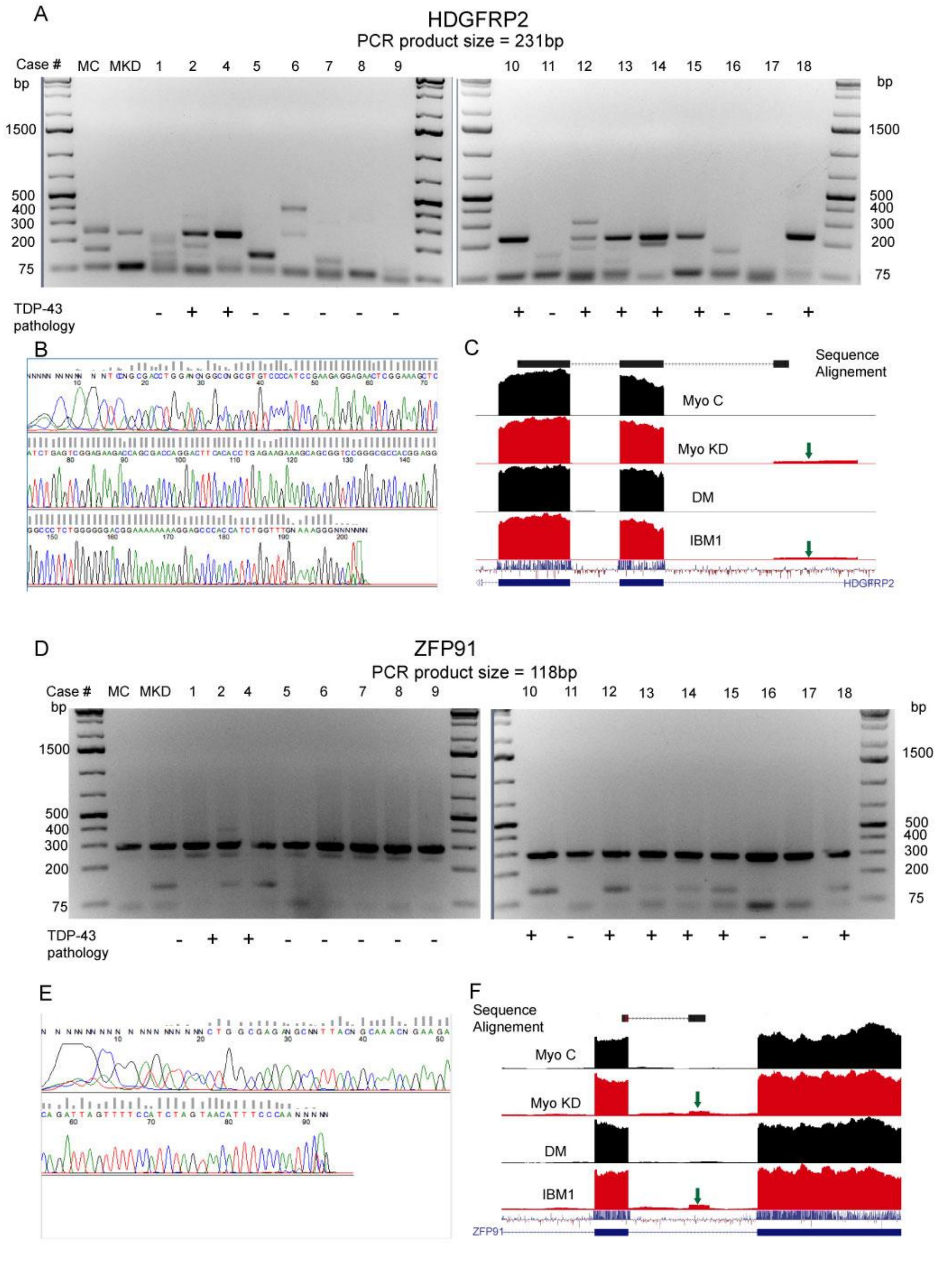

**Fig. S3. Sequence and alignment validation of cryptic exon RT-PCR: HDGFRP2/ZFP91**

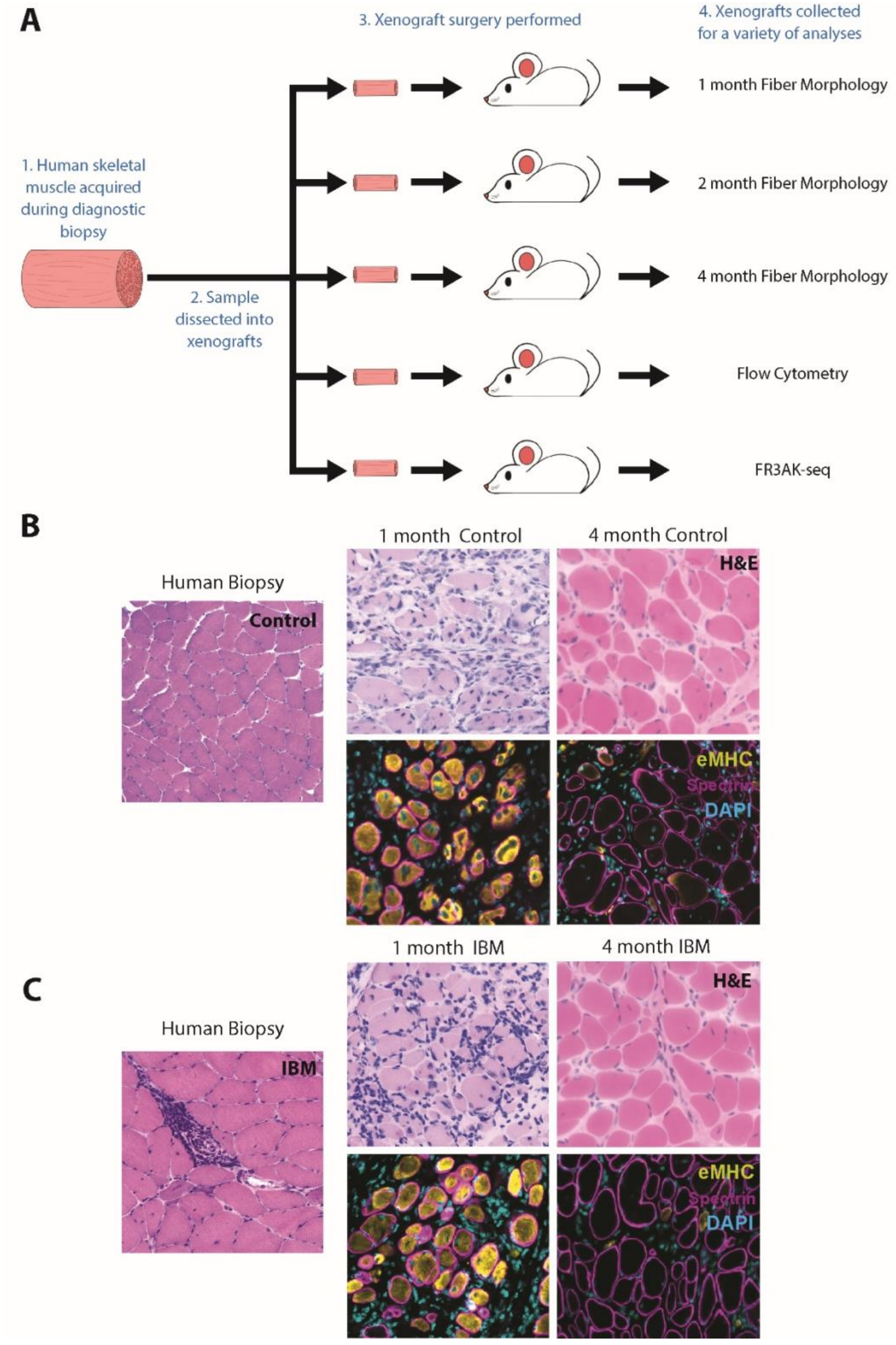

**Fig. S4. Description of xenograft model.** (A) Human skeletal muscle biopsy specimens are obtained from both IBM and control patients, dissected into ~7 x 3 x 3mm xenografts, and transplanted into NOD-Rag1$^{null}$IL2r$\gamma^{null}$ (NRG) mice which lack the ability to generate mature B or T cells. On average, 8 xenografts can be generated from a single patient biopsy specimen. These xenografts are then harvested at a variety of different timepoints for various analyses, such as fiber morphology analysis, flow cytometry, and T cell receptor sequencing via FR3AK-seq. Due to the small size of the xenograft, the xenograft must be harvested in its entirety, and therefore, any individual xenograft can only be used for a single timepoint. (B,C) Xenografts mature over a 4-month period as shown by representative images of 1-month and 4-month control (B) and IBM (C) xenografts. Following the patient biopsy (representative H&E shown for a control (B) and IBM (C) patient biopsy), all mature myofibers undergo degeneration within 1-2 weeks, and satellite cells proliferate and differentiate into myoblasts which then fuse to form newly regenerated myofibers by 1 month (labeled with embryonic myosin (eMHC, yellow) (Zhang et al, 2014). As xenografts mature, the fibers enlarge and expression of eMHC decreases over time. Human-specific anti-spectrin antibodies label the myofiber sarcolemmal membrane (magenta), and DAPI (blue) stains all nuclei. For video of xenograft procedure and detailed protocol, see Britson et al, JOVE, 2019.

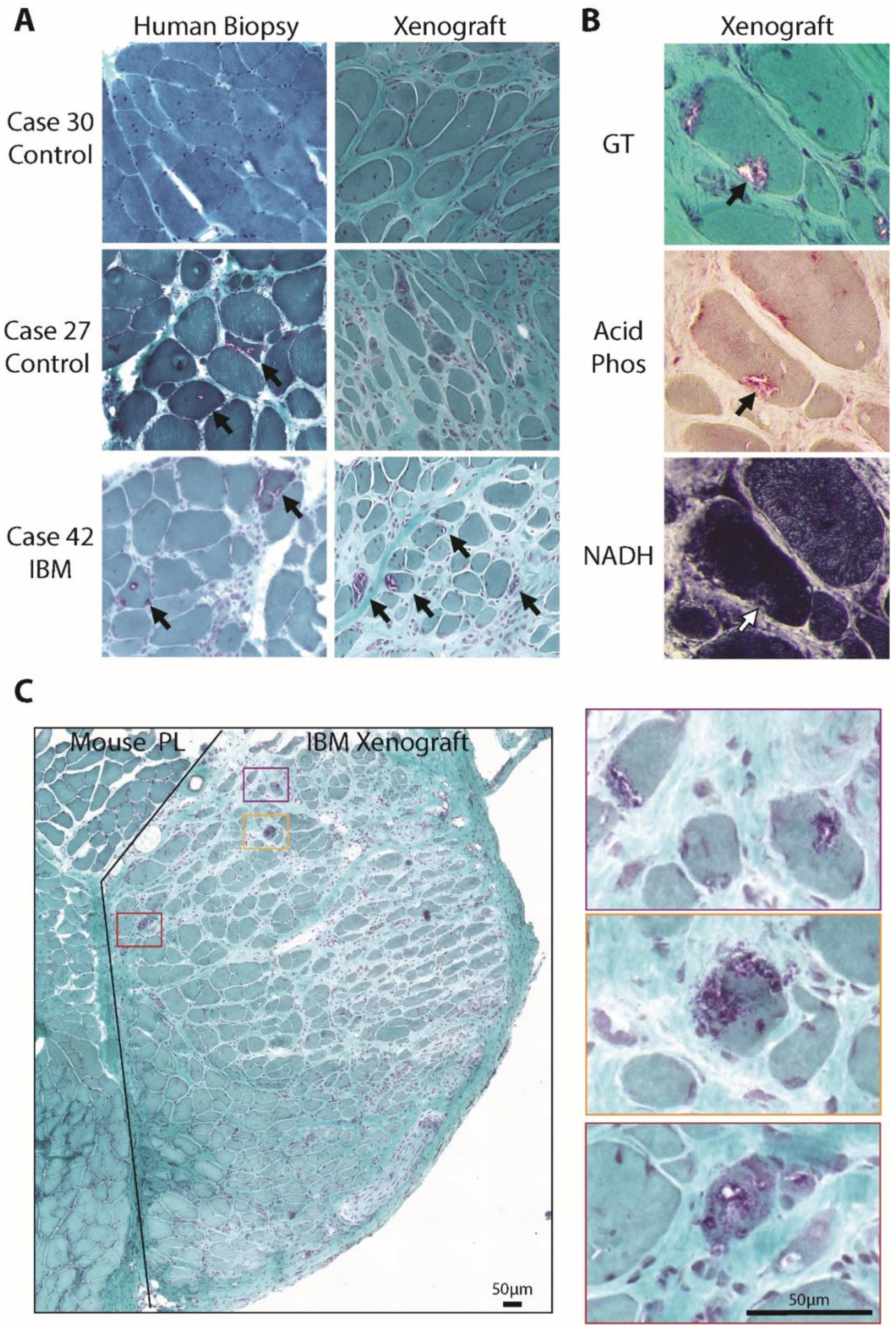

**Fig. S5. Rimmed vacuoles in the xenograft model.** (A) Representative images of Gomori trichrome stains in human biopsies and 4-month xenografts from cases 30, 27, and 42. The Case

27 patient was diagnosed with a genetic vacuolar myopathy and clear examples of rimmed vacuoles can be seen in both the patient biopsy and 4-month xenograft (arrows). Rimmed vacuoles are also observed in IBM patient biopsies and 4-month IBM xenografts (arrows). (B) Representative images of Gomori trichrome (GT), Acid Phosphatase (Acid Phos), and NADH staining of a myofiber containing a rimmed vacuoles (arrow) from a 4-month IBM xenograft demonstrating typical features of autophagolysosomes including acid phosphatase positivity and light NADH staining. (C) A representative image of a 4-month IBM xenograft with adjacent mouse peroneus longus (PL) muscle stained with Gormori Trichrome. Inserts (purple, orange, red) show increased magnifications of examples of rimmed vacuoles. Scale bars show 50µm.

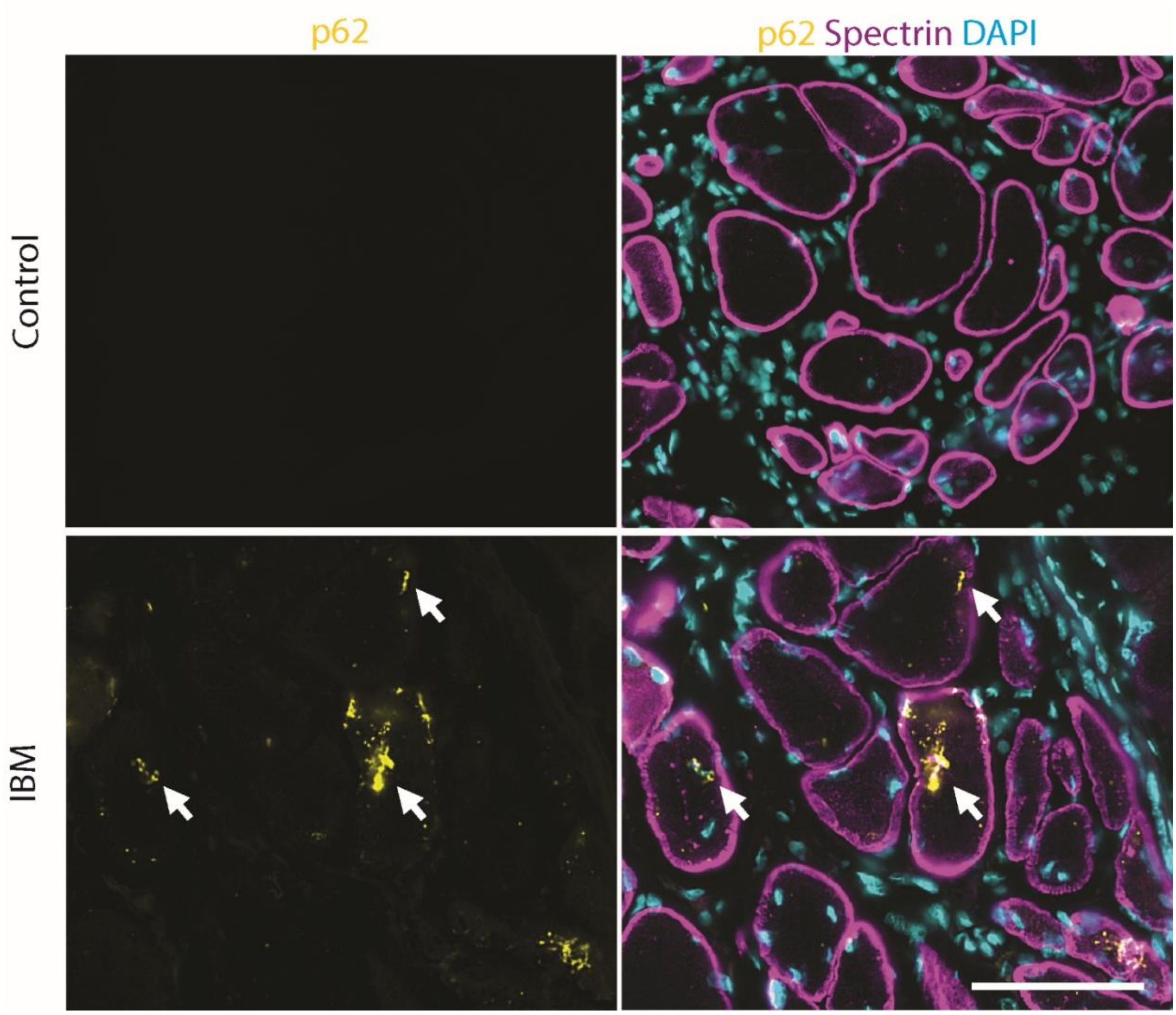

**Fig. S6. IBM xenografts show p62 aggregation at 4 months.** Representative images of autophagic adaptor protein p62 staining within 4-month control and IBM xenografts. Fibers in IBM xenografts with p62 positive aggregates are indicated with arrows. Scale bar shows 50μm.

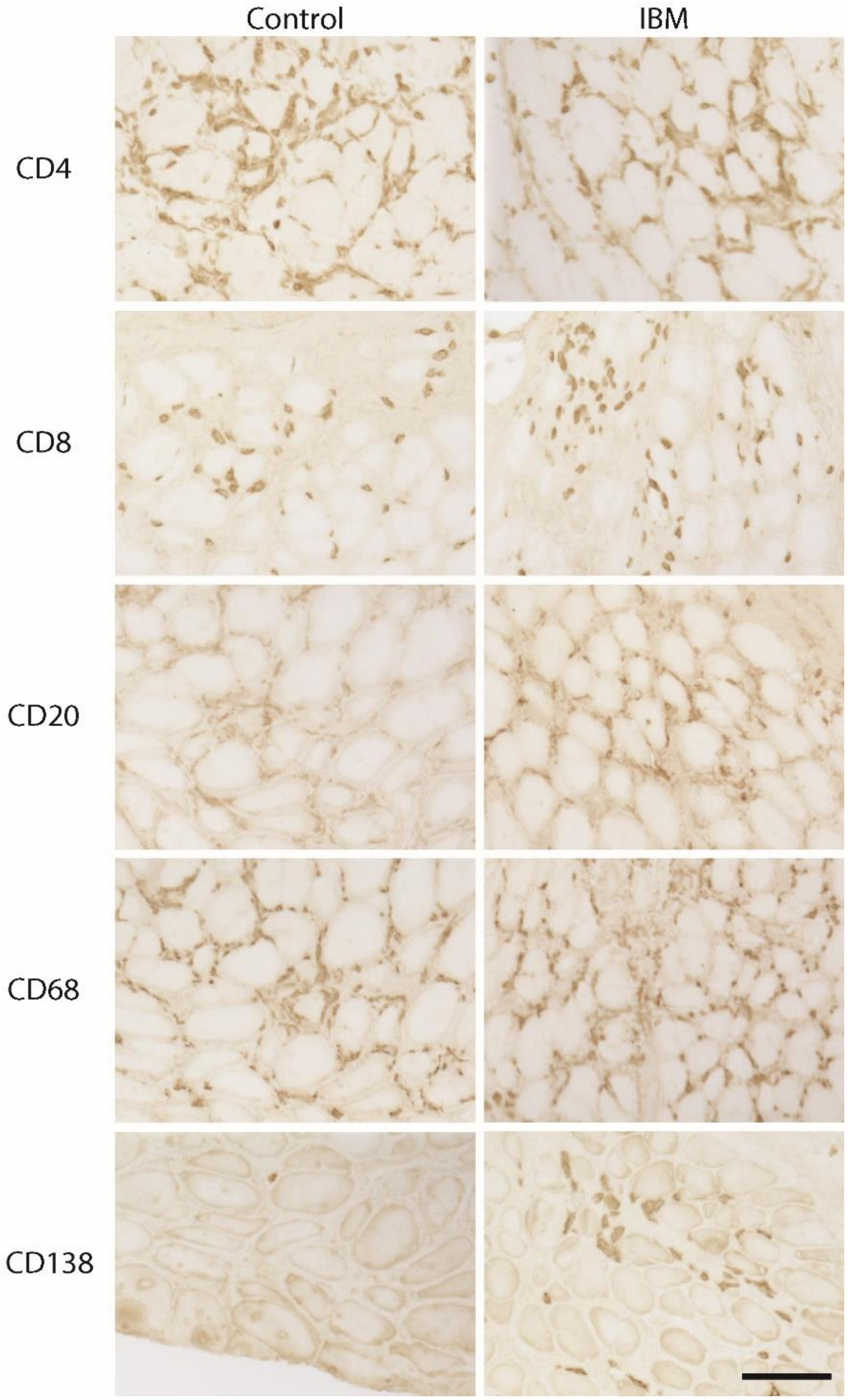

**Fig. S7. Multiple human immune cell types are present within xenografts.**

Representative immunostains of CD4 (helper T cells), CD8 (cytotoxic T cells), CD20 (B cells), CD68 (macrophages), and CD138 (plasma cells) from 4-month xenografts are shown. Scale bar shows 100µm.

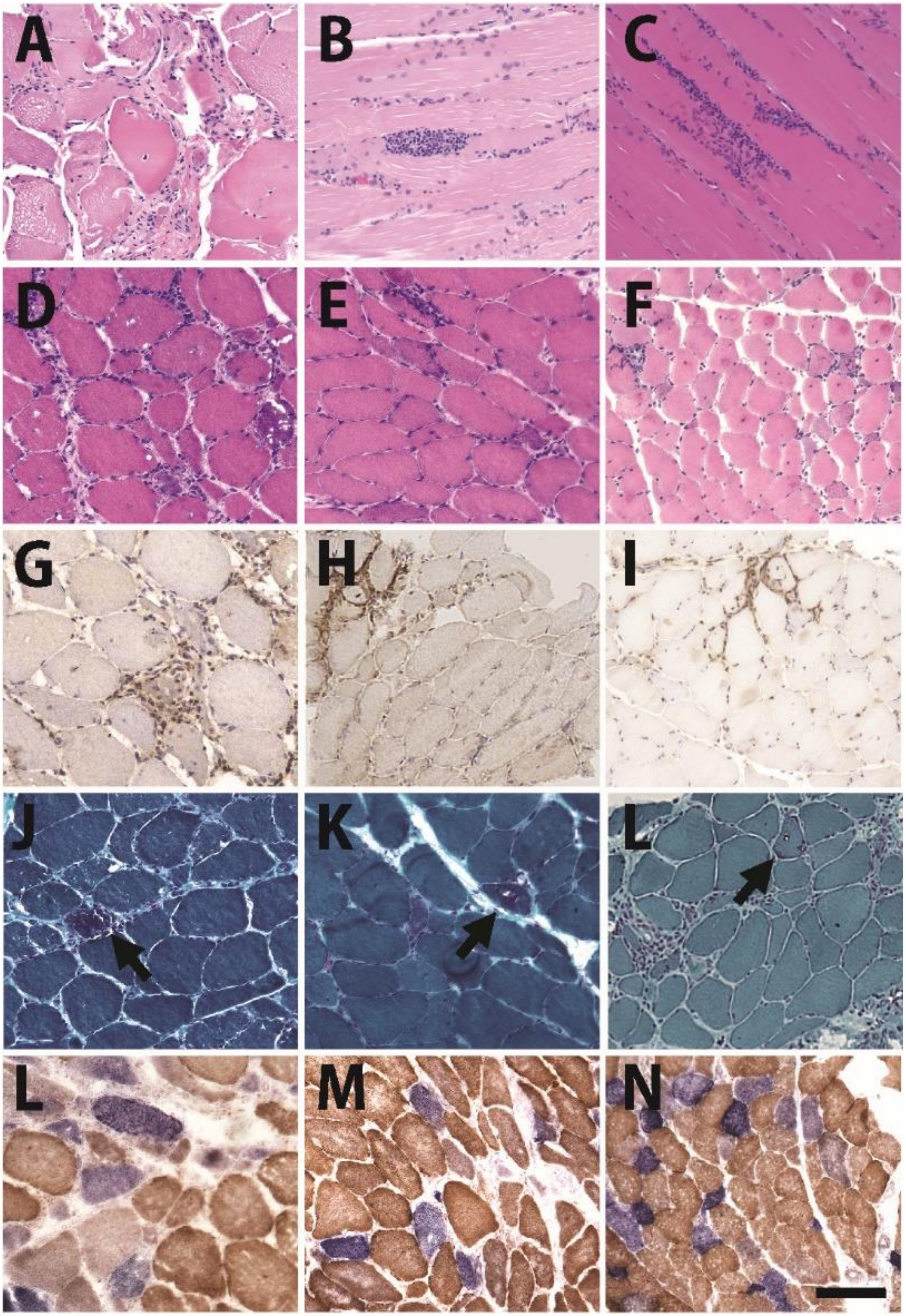

**Fig. S8: Histological features of human biopsies in OKT3 experiments.** Paraffin (A-C), H&E

(D-F), CD3 (G-I), Gomori Trichrome (GT) (J-L), and dual COX-SDH (M-O) stains showing representative histology from IBM cases 23, 26, And 36. All biopsies show endomysial inflammation and primary invasion (A-I). In addition, all biopsies show many COX deficient fibers (L-M). The IBM cases 23, 26, and 42 show rimmed vacuoles (J, K arrows), but vacuoles are absent in IBM case 36. Scale bar shows 100μm.

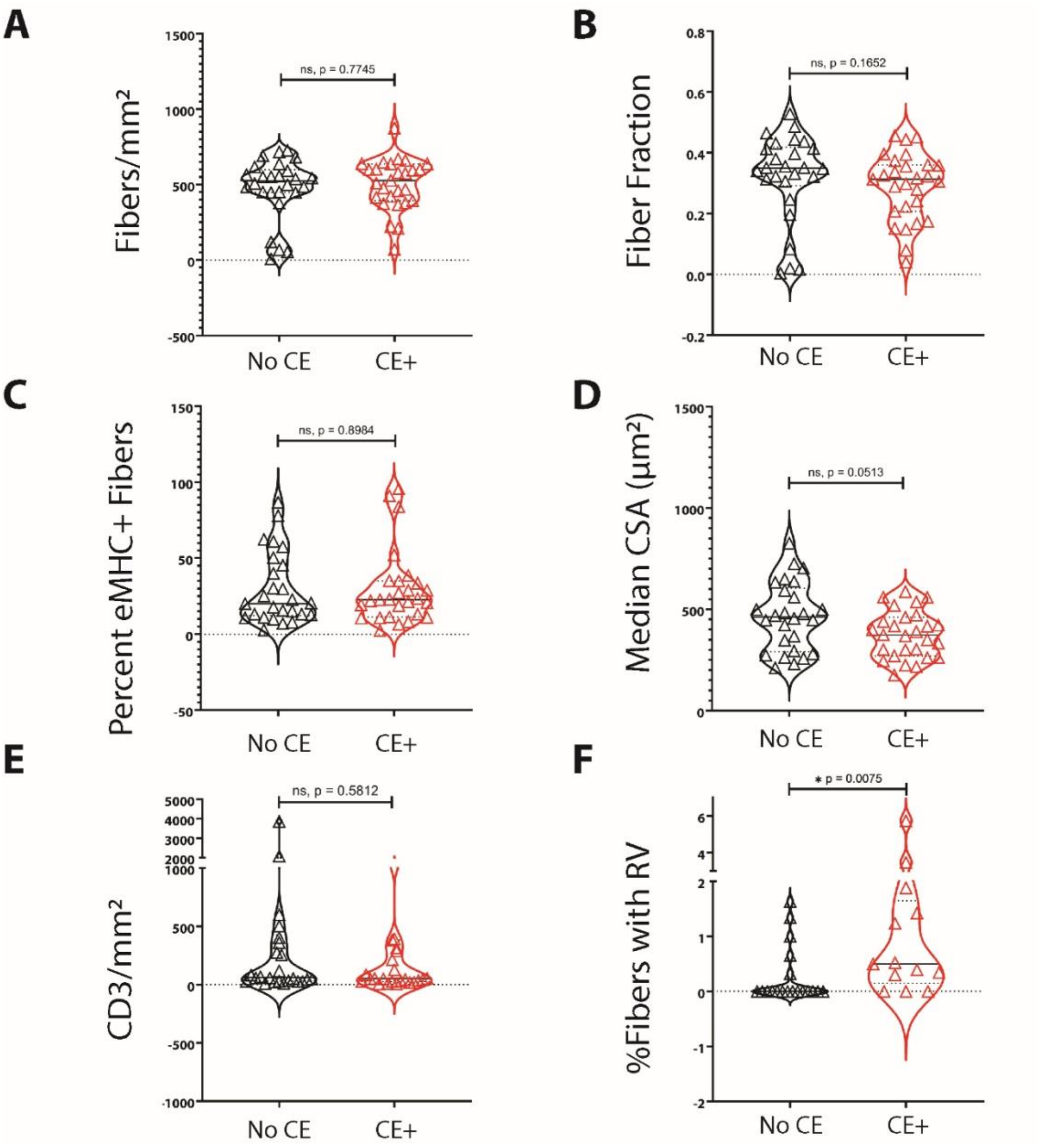

**Fig. S9. Rimmed vacuoles are more frequent in xenografts containing cryptic exons.** (A-F) Comparisons of xenografts with (CE+) or without (No CE) cryptic exon expression. For all comparisons, No CE: n = 26 xenografts; CE+; n = 27 xenografts except for the %Fibers with rimmed vacuoles (RV) where No CE; n = 19 and CE+; n = 13. Mann-Whitney test was used to determine significance (*p ≤ 0.05).

| Gene Symbol | UCSC ID | Cryptic Exon Location (hg19) | Verified Length (bp) | Present in IBM | Present in HeLa | Myoblast Cryptic (%) | Inf. Ins. | 5' UTR or ncRNA | Start Site | PolyA | Exon Ext. | NMD | Myoblast Control | Myoblast TDP-43 Knockdown | Control vs. Knockdown (FoldChange) |
|---|---|---|---|---|---|---|---|---|---|---|---|---|---|---|---|
| ACOT11 | uc001cxl.2 | chr1:55,066,636-55,066,745 | 110 | *** | Y | 89 | -- | -- | -- | -- | -- | Y | 8.3 | 0.6 | -12.8 |
| ZNF529 | uc002oeh.4 | chr19:37,063,945-37,064,041 | 97 | N | Y | 64 | -- | 5'UTR | -- | -- | Y | -- | 2.2 | 2.7 | 1.2 |
| CDO1 | uc003krg.3 | chr5:115,149,595-115,149,651 | 57 | N | N | 64 | Y | -- | -- | -- | -- | -- | 1.9 | 1.3 | -1.4 |
| CEP72 | uc003jbf.3 | chr5:648,215-648,338 | 124 | N | Y | 49 | -- | -- | -- | -- | -- | Y | 0.8 | 0.9 | 1.1 |
| PPP2R2D | uc001lks.3 | chr10:133,770,158-133,773,339 | 3,182 | N | Y | 32 | -- | -- | -- | Y | -- | -- | 17.6 | 11.1 | -1.6 |
| USP13 | uc003fkh.3 | chr3:179,418,918-179,418,967 | 50 | N | Y | 30 | -- | -- | -- | -- | -- | Y | 4 | 1.7 | -2.4 |
| SLC17A9 | uc002yea.4 | chr20:61,588,933-61,591,565 | 2,633 | N | N | 30 | -- | -- | -- | -- | Y | Y | 1.4 | 2.3 | 1.6 |
| RHEBL1 | uc001rtc.1 | chr12:49,463,375-49,463,516 | 142 | N | N | 27 | -- | -- | -- | -- | Y | Y | 1.1 | 4.8 | 4.5 |
| ZC3H12C | uc010rwc.2 | chr11:109,959,110-109,959,327 | 218 | N | N | 24 | -- | -- | Y | -- | -- | -- | 3.8 | 6.3 | 1.6 |
| SLC39A8 | uc003hwc.2 | chr4:103,226,269-103,227,158 | 890 | N | N | 19 | -- | -- | -- | -- | Y | Y | 2.7 | 6.5 | 2.4 |
| ADAMTS6 | uc003jtp.3 | chr5:64,605,352-64,605,408 | 57 | N | N | 19 | Y | -- | -- | -- | -- | -- | 2.7 | 2.8 | 1.1 |
| ST5 | uc001mgt.3 | chr11:8,801,680-8,801,798 | 119 | N | Y | 14 | -- | 5'UTR | -- | -- | -- | -- | 6.7 | 10.1 | 1.5 |
| FAM114A2 | uc003lvc.3 | chr5:153,416,928-153,417,015 | 88 | N | Y | 14 | -- | 5'UTR | -- | -- | -- | -- | 18.2 | 10.8 | -1.7 |
| BIRC3 | uc001pgx.3 | chr11:102,190,608-102,190,737 | 130 | Y | N | 13 | -- | 5'UTR | -- | -- | -- | -- | 1.8 | 7.4 | 4.1 |
| PFKP | uc001igp.3 | chr10:3,141,749-3,142,011 | 263 | N | Y | 13 | -- | -- | -- | -- | -- | Y | 73.5 | 42 | -1.8 |
| ACSF2 | uc002iqu.2 | chr17:48,539,181-48,539,246 | 66 | Y | Y | 12 | Y | -- | -- | -- | -- | -- | 4.9 | 5.3 | 1.1 |
| XPO4 | uc001unq.4 | chr13:21,374,252-21,374,294 | 43 | N | Y | 12 | -- | -- | -- | -- | Y | Y | 5.4 | 5.3 | -1 |
| HDGFRP2 | uc002mao.3 | chr19:4,492,012-4,492,149 | 138 | Y | Y | 11 | Y | -- | -- | -- | -- | -- | 17.5 | 18.5 | 1.1 |
| COL4A5 | uc004enz.2 | chrX:107,836,061-107,836,104 | 44 | N | N | 10 | -- | -- | -- | -- | -- | Y | 15.4 | 7.9 | -1.9 |
| ZFP91 | uc001nmx.4 | chr11:58,384,466-58,384,527 | 62 | Y | Y | 8 | -- | -- | -- | -- | -- | N | 25.9 | 22.2 | -1.2 |
| GPSM2 | uc010ovc.2 | chr1:109,438,939-109,439,053 | 115 | Y | Y | 8 | -- | -- | -- | -- | -- | Y | 19 | 8.8 | -2.1 |
| PKN1 | uc002myp.3 | chr19:14,560,900-14,561,129 | 230 | N | Y | 7 | -- | -- | -- | -- | -- | Y | 18.3 | 9.5 | -1.9 |
| RANBP1 | uc002zro.1 | chr22:20,110,103-20,110,220 | 118 | N | Y | 6 | -- | -- | -- | -- | -- | Y | 63.6 | 33.3 | -1.9 |
| BCL2L13 | uc002zmw.4 | chr22:18,205,040-18,205,140 | 101 | Y | N | 5 | -- | -- | -- | -- | -- | N | 17.4 | 12.1 | -1.4 |

**Table S1. TDP-43 associated cryptic exons in human myoblasts.**

Table showing cryptic exons annotated by genomic location and size (bp).

| Patient | Patient Sex | Age at biopsy | Diagnosis | CK | Biopsy | EMG | Atrophy | RV | inflam | mito |
|---|---|---|---|---|---|---|---|---|---|---|
| 1 | female | 63 | DM | 1614 | rectus | Nonirritable myopathy | perifascicular | No | none | No |
| 2 | male | 72 | IBM | 142 | rectus | Irritable myopathy | yes | Yes | primary | Yes |
| 4 | male | 72 | IBM | 1986 | rectus | Irritable myopathy | yes | No | primary | Yes |
| 5 | male | 65 | denerv | 1039 | rectus | normal | neurogenic | No | none | No |
| 6 | female | 56 | normal | 97 | rectus | neurogenic | no | No | none | No |
| 7 | male | 70 | DM | 33 | rectus | Irritable myopathy | minimal | No | none | No |
| 8 | male | 68 | DM | 1164 | rectus | Irritable myopathy | type 2 | No | none | No |
| 9 | male | 57 | DM | 341 | rectus | Nonirritable myopathy | perifascicular | No | perivasc | No |
| 10 | male | 54 | IBM | 1568 | rectus | Irritable myopathy | severe | Yes | primary | Yes |
| 11 | male | 67 | denerv | 571 | rectus | neurogenic | neurogenic | No | None | No |
| 12 | female | 66 | IBM | 200 | rectus | neurogenic | yes | Yes | primary | Yes |
| 13 | male | 46 | IBM | 336 | rectus | neurogenic | yes | No | primary | Yes |
| 14 | male | 64 | IBM | 180 | rectus | Irritable myopathy | severe | Yes | primary | Yes |
| 15 | male | 64 | IBM | 939 | rectus | Irritable myopathy | yes | Yes | primary | Yes |
| 16 | male | 61 | normal | 119 | rectus | normal | minimal | No | none | No |
| 17 | male | 61 | normal | 1628 | rectus | normal | yes | No | none | No |

**Table S2. Demographics and clinical and muscle biopsy findings in Cryptic Exon Patient Cohort.**

Age at muscle biopsy (average: IBM 62 yrs; control 63yrs), and sex ratio (average: IBM 85.7% male; control 77.8% male) are similar between IBM cases and controls. All IBM cases met ENMC 2011 criteria for clinically defined IBM, and Dermatomyositis (DM) cases meet Bohan and Peter criteria. Serum Creatine Kinase (CK) measured in international units/Liter at time of muscle biopsy is shown. All biopsies were of rectus femoris muscle. “denerv” in diagnosis column refers to findings of denervation, or “neurogenic atrophy” on muscle biopsy. “EMG” refers to primary pattern of changes observed on electromyography study performed prior to muscle biopsy; “nonirritable” denotes normal spontaneous activity whereas “irritable” refers to abnormal spontaneous activity with positive waves and/or fibrillations. “Neurogenic” EMG refers to an EMG that shows reduced recruitment and motor unit action potentials with prolonged amplitude and/or duration. RV: rimmed vacuoles; inflam: inflammation; perivasc: perivascular. Primary inflammation refers to presence of endomysial inflammatory cells surrounding and invading normal appearing myofibers. “mito” refers to excessive number of COX-negative fibers suggesting mitochondrial abnormalities.

**Cryptic Exon Patient Cohort**

| Patient | Sample | Group | GP | HDG | ACSF | ZFP |
|---|---|---|---|---|---|---|
| 1 | 2005-66 | DM | | | | |
| 2 | 2006-246 | IBM | ✓ | ✓ | ✓ | ✓ |
| 4 | 2007-76 | IBM | ✓ | ✓ | ✓ | ✓ |
| 5 | 2008-112 | neurogenic | | | | |
| 6 | 2008-173 | Normal | | | | |
| 7 | 2009-349 | DM | | | | |
| 8 | 2010-127 | DM | | | | |
| 9 | 2011-257 | DM | | | | |
| 10 | 2011-375 | IBM | ✓ | ✓ | ✓ | ✓ |
| 11 | 2011-377 | neurogenic | | | | |
| 12 | 2012-75 | IBM | ✓ | ✓ | ✓ | ✓ |
| 13 | 2013-10 | IBM | ✓ | ✓ | ✓ | ✓ |
| 14 | 2013-181 | IBM | ✓ | ✓ | ✓ | ✓ |
| 15 | 2013-339 | IBM | ✓ | ✓ | ✓ | ✓ |
| 16 | 2013-342 | Normal | | | | |
| 17 | 2013-408 | Normal | | | | |
| A | 2003-194 | Anti-HMGCR | | | | |
| B | 2004-254 | Anti-Jo-1 | | | | |
| C | 2006-122 | SSc | | | | |
| D | 2007-184 | SSc | | | | |
| E | 2008-118 | Anti-MDA5 | | | | |
| F | 2008-127 | Anti-PL-12 | | | | |
| G | 2008-140 | Anti-Jo-1 | | | | |
| H | 2008-212 | Anti-HMGCR | | | | |
| I | 2008-292 | Anti-HMGCR | | | | |
| J | 2008-91 | Anti-HMGCR | | | | |
| K | 2008-97 | Anti-HMGCR | | | | |
| L | 2009-141 | IBM | - | | | |
| M | 2009-174 | SSc | | | | |
| N | 2009-187 | SSc | | | | |
| O | 2009-212 | Anti-Jo-1 | | | | |
| P | 2009-347 | SSc | | | | |
| Q | 2009-386 | Anti-PL-7 | | | | |
| 18 | 2009-410 | SSc | | | | |
| 19 | 2009-57 | IBM | ✓ | ✓ | ✓ | |
| 20 | 2010-105 | Anti-Mi-2 | | | | |
| 21 | 2010-148 | Anti-TIF1g | | | | |
| 22 | 2010-221 | SSc | | | | |
| 23 | 2010-31 | Anti-HMGCR | | | | |
| 24 | 2010-436 | Anti-SRP | | | | |
| 25 | 2010-7 | IBM | ✓ | ✓ | ✓ | ✓ |
| 26 | 2011-120 | IBM | | ✓ | | |
| 27 | 2011-189 | SSc | | | | |
| 28 | 2011-216 | Anti-HMGCR | | | | |
| 29 | 2011-25 | Anti-HMGCR | | | | |
| 30 | 2011-381 | Anti-SRP | | | | |
| 31 | 2011-64 | SSc | | | | |

| Patient | Sample | Group | GP | HDG | ACSF | ZFP |
|---|---|---|---|---|---|---|
| 32 | 2011-65 | SSc | | | | |
| 33 | 2012-191 | IBM | ✓ | ✓ | ✓ | ✓ |
| 34 | 2012-271 | Anti-Jo-1 | | | | |
| 35 | 2013-178 | Anti-HMGCR | | | | |
| 36 | 2013-208 | IBM | ✓ | ✓ | ✓ | ✓ |
| 37 | 2013-242 | Anti-HMGCR | | | | |
| 38 | 2013-319 | Anti-HMGCR | | | | |
| 39 | 2013-409 | Anti-SRP | | | | |
| 40 | 2013-411 | Anti-SRP | | | | |
| 41 | 2013-426 | Anti-NXP2 | | | | |
| 42 | 2013-432 | IBM | | | | |
| 43 | 2013-88 | Anti-HMGCR | | | | |
| 44 | 2014-277 | Anti-PM/SCL | | | | |
| 45 | 2014-306 | SSc | | | | |
| 46 | 2014-72 | Anti-Jo-1 | | | | |
| 47 | 2014-81 | Anti-HMGCR | | | | |
| 48 | 2015-203 | Normal | | | | |
| 49 | 2015-206 | SSc | | | | |
| 50 | 2015-207 | Normal | | | | |
| 51 | 2015-247 | Normal | | | | |
| 52 | 2015-272 | Normal | | | | |
| 53 | 2015-294 | Normal | | | | |
| 54 | 2015-295 | SSc | | | | |
| 55 | 2015-305 | Normal | | | | |
| 56 | 2015-306 | Normal | | | | |
| 57 | 2015-339 | Normal | | | | |
| 58 | 2015-343 | Normal | | | | |
| 59 | 2015-418 | Normal | | | | |
| 60 | 2014-166 | IBM | ✓ | | ✓ | |
| 61 | 2014-185 | IBM | | | | |
| 62 | 2014-215 | IBM | ✓ | | ✓ | |
| 63 | 2014-344 | IBM | ✓ | | ✓ | |
| 64 | 2015-195 | IBM | ✓ | | ✓ | |
| 65 | 2015-279 | IBM | ✓ | | ✓ | |
| 66 | 2015-367 | IBM | | | ✓ | |
| 67 | 2015-408 | IBM | ✓ | | | |
| 68 | 2016-77 | IBM | ✓ | | ✓ | |
| 69 | 2016-92 | IBM | ✓ | | ✓ | |
| 70 | 2016-194 | IBM | ✓ | | ✓ | |
| 71 | 2016-195 | IBM | | | ✓ | |
| 72 | 2017-300 | IBM | ✓ | | ✓ | |
| 73 | 2018-330 | anti-MJ | | | | |

| | |
|---|---|
| SSc | scleroderma and systemic sclerosis |
| IBM | inclusion body myositis |
| Normal | |

**Xenograft Patient Cohort**

| Case | Sample | Group | GP | HDG | ACSF | ZFP |
|---|---|---|---|---|---|---|
| 1 | 2015-334 | IBM | ✓ | ✓ | ✓ | |
| 2 | 2016-157 | IBM | | ✓ | | |
| 3 | 2017-175 | Control | | | | |
| 4 | 2017-189 | Control | | | | |
| 5 | 2017-282 | IBM | ✓ | ✓ | | |
| 6 | 2017-330 | IBM | ✓ | ✓ | ✓ | |
| 8 | 2018-87 | IBM | ✓ | ✓ | ✓ | |
| 9 | 2018-126 | IBM | | | | |
| 10 | 2018-128 | IBM | | ✓ | | |
| 13 | 2018-193 | IBM | ✓ | ✓ | ✓ | |
| 15 | 2018-226 | Control | | | | |
| 19 | 2018-280 | IBM | ✓ | ✓ | ✓ | |
| 20 | 2018-303 | IBM | | | | |
| 21 | 2018-309 | IBM | ✓ | | | |
| 22 | 2019-7 | Control | | | | |

| Case | Sample | Group | GP | HDG | ACSF | ZFP |
|---|---|---|---|---|---|---|
| 23 | 2019-36 | IBM | | | | |
| 26 | 2019-86 | IBM | ✓ | ✓ | | |
| 27 | 2019-101 | Control | ✓ | ✓ | | |
| 29 | 2019-120 | Control | | | | |
| 30 | 2019-138 | Control | | | | |
| 31 | 2019-153 | Control | | | | |
| 33 | 2019-171 | IBM | | | | |
| 34 | 2019-215 | Control | | | | |
| 35 | 2019-228 | Control | | | | |
| 36 | 2019-231 | IBM | | ✓ | | |
| 38 | 2019-290 | Control | | | | |
| 40 | 2019-311 | IBM | | | | |
| 41 | 2019-312 | Control | | | | |
| 42 | 2019-322 | IBM | ✓ | ✓ | ✓ | |

**Table S3. Patient cohort cryptic exon expression.** A summary of the cryptic exon expression for the Cryptic exon and Xenograft patient cohorts. Out of 119 total patients, 36/44 IBM patients and only 1/74 control patients were positive for cryptic exons, resulting in a sensitivity of 82% and specificity of 99% for diagnosis of IBM. In addition, this difference was found to be significant by Fisher's Exact Test (IBM, $p<0.0001$). A checkmark indicates cryptic exon expression, and a grey box indicates a cryptic exon target whose expression was not tested.

| | Controls | | | | | | | | | | | | |
|---|---|---|---|---|---|---|---|---|---|---|---|---|---|
| Case | Patient Sex | Age at biopsy | Duration (yrs) | CK | Biopsy | Diagnosis | ENMC crit | FF > SA | KE >= HF | Inflam | invasion | RV | Increased COX neg fibers | MHC-I |
| 3 | Male | 57 | 1 | 67 | Deltoid | Dermatomyositis | no | no | no | yes | no | no | no | yes (perifasc) |
| 4 | Male | 71 | 14 | 306 | Biceps | myalgia, cramps | no | no | no | no | no | no | no | no |
| 15 | Female | 64 | 9 | 92 | Rectus | metabolic myopathy | no | no | no | no | no | no | no | no |
| 18 | Female | 39 | 2 | 65 | Rectus | Dermatomyositis | no | no | no | no | no | no | no | no |
| 22 | Male | 62 | 4 | 266 | Vastus | neurogenic atrophy | no | no | yes | no | no | no | no | no |
| 27 | Female | 60 | 3 | 5,408 | Biceps | Vacuolar Myopathy | no | no | no | no | no | many | no | NL |
| 29 | Male | 55 | 2 | 449 | Biceps | Metabolic Myopathy | no | no | no | no | no | no | no | no |
| 30 | Male | 68 | 1 | 224 | Rectus | inflamm myopathy | no | no | no | perivasc | no | no | no | incr |
| 31 | Female | 50 | <1 | 51 | Rectus | Inflamm myopathy | no | no | no | rare | no | no | no | NL |
| 34 | Female | 76 | 9 | 144 | Biceps | mitochondrial myopathy | no | no | no | no | no | no | yes | no |
| 35 | Male | 68 | 4 | 219 | Biceps | normal | no | no | no | no | no | no | no | no |
| 38 | Male | 65 | 5 | 362 | Biceps | neurogenic atrophy | no | no | no | perivasc | no | no | no | NL |
| | sIBM | | | | | | | | | | | | | |
| Case | Patient Sex | Age at biopsy | Duration (yrs) | CK | Biopsy | Diagnosis | ENMC crit | FF > SA | KE >= HF | Inflam | invasion | RV | Increased COX neg fibers | MHCI |
| 5 | Male | 77 | 3 | 198 | Bicep | sIBM | CD | yes | yes | yes | yes | yes | yes | incr |
| 6 | Male | 74 | 7 | 303 | Vastus | sIBM | Prob | no | yes | yes | yes | yes | yes | incr |
| 8 | Male | 68 | 1 | 198 | Bicep | sIBM | CD | yes | yes | yes | no | rare | no | incr |
| 9 | Male | 77 | 8 | 1700 | Bicep | sIBM | Prob | yes | no | yes | yes | no | yes | incr |
| 10 | Male | 53 | 2 | 339 | Deltoid | sIBM | CD | yes | yes | yes | yes | yes | no | min incr |
| 13 | Male | 67 | 3 | 584 | Bicep | sIBM | CD | yes | yes | yes | yes | yes | no | incr |
| 19 | Female | 66 | 3 | 800 | Bicep | sIBM | CD | yes | yes | yes | yes | yes | yes | |
| 20 | Male | 66 | 2 | 503 | Bicep | sIBM | Prob | yes | no | yes | yes | yes | yes | |
| 21 | Female | 74 | 2 | 820 | Deltoid | sIBM | CD | yes | yes | yes | yes | no | yes | incr |
| 23 | Male | 64 | 1 | 1400 | Bicep | sIBM | Prob | yes | no | yes | yes | yes | yes | incr |
| 26 | Male | 75 | 5 | 629 | Bicep | sIBM | CD | yes | yes | yes | yes | rare | yes | incr |
| 33 | Male | 80 | 7 | 205 | Vastus | sIBM | Prob | yes | no | yes | no | yes | no | mild incr |
| 36 | Male | 64 | 4 | 210 | Rectus | sIBM | Prob | yes | no | yes | yes | no | yes | NL |
| 40 | Female | 57 | 1 | 468 | Biceps | sIBM | Prob | yes | no | yes | yes | no | no | incr |
| 42 | Male | 77 | 3 | 469 | Vastus | sIBM | Prob | yes | no | yes | yes | yes | no | incr |

**Table S4**. **Clinical, demographic, and biopsy details of patients in the Xenograft Cohort of this study.** The disease duration was determined from the time of symptom onset to the time the diagnostic biopsy was performed. Serum Creatine Kinase (CK) measured in international units/Liter at time of muscle biopsy is shown. Diagnosis of controls refers to working clinical diagnosis following muscle biopsy. All IBM cases met ENMC 2011 criteria (ENMC crit) for clinically defined (CD) or probable (Prob) IBM ([4]). FF: finger flexion; SA: shoulder abduction KE: knee extension; HF: hip flexion; inflam: inflammation; perivasc: perivascular; RV: rimmed vacuoles; "Increased Cox neg fibers" refers to excessive number of COX-negative fibers suggesting mitochondrial abnormalities. Invasion refers to presence of endomysial inflammatory cells surrounding and invading normal appearing myofibers. MHC-I overexpression is indicated at a variety of different levels: yes; no; NL: normal; incr: increased; min incr: minimally increased; mild incr: mild increase).

| | Entire Population (n=27) | Controls (n=12) | IBM (n=15) | p-value controls vs IBM |
|---|---|---|---|---|
| **Sex (male), n (%)** | 19 (70.37%) | 7 (58.3%) | 12 (85.7%) | 0.3981 |
| **Age (yrs), mean ± SD** | 65.7 ± 9.62 | 61.2 ± 10.03 | 70.1 + 7.39 | 0.0373 |
| **Disease duration (yrs), mean ± SD** | 4.02 ± 3.25 | 4.77 ± 4.24 | 3.64 ± 2.27 | 0.5973 |
| **Biopsy location, n (%)** | | | | |
| **- Biceps** | 13 (48.15%) | 5 (41.67%) | 8 (57.1%) | |
| **- Rectus femoris** | 4 (14.81%) | 2 (16.67%) | 2 (14.3%) | |
| **- Vastus lateralis** | 4 (14.81%) | 2 (16.67%) | 2 (14.3%) | |
| **- Deltoid** | 2 (7.40%) | 0 (0%) | 2 (14.3%) | |

**Table S5. Summary of the sex ratio, age, disease duration and biopsy location of the xenograft patient Cohort.** The disease duration was determined from the time of symptom onset to the time the diagnostic biopsy was performed. Mann-Whitney test was used to determine significance for the following features: age and disease duration. Fisher’s exact test was used to determine significance for the sex ratio of the patient population.

| Xenograft | Group | Timepoint | GPSM | HDGF | ACSF |
|---|---|---|---|---|---|
| 3.5 | Control | 8 month | | | |
| 3.7 | Control | 8 month | | | |
| 3.2 | Control | 10 month | | | |
| 3.6 | Control | 10 month | | | |
| 4.4 | Control | 4 month | | | |
| 4.5 | Control | 8 month | | | |
| 4.6 | Control | 8 month | | | |
| 4.3 | Control | 10 month | | | |
| 4.7 | Control | 10 month | | | |
| 15.2 | Control | 4 month | | | |
| 15.5 | Control | 4 month | | | |
| 15.4 | Control | 8 month | | | |
| 18.2R | Control | 4 month | | | |
| 22.2L | Control | 4 month | | | |
| 22.6R | Control | 8 month | | | |
| 22.6L | Control | 8 month | | | |
| 22.8R | Control | 10 month | | | |
| 22.8L | Control | 10 month | | | |
| 27.1 | Control | 4 month | | | |
| 29.1 | Control | 4 month | | | |
| 31.8R | Control | 8 month | | | |
| 31.8L | Control | 8 month | | | |
| 31.7R | Control | 10 month | | | |
| 31.7L | Control | 10 month | | | |
| 35.3 | Control | 8 month | | | |
| 35.4 | Control | 8 month | | | |
| 35.5 | Control | 8 month | | | |
| 38.3R | Control | 8 month | | | ✓ |
| 38.3L | Control | 8 month | | | |
| 41.7R | Control | 8 month | | | |
| 41.7L | Control | 8 month | | | |
| 41.4 | Control | 10 month | | | |
| 41.9R | Control | 10 month | | | |
| 41.9L | Control | 10 month | | | |
| 5.5L | IBM | 4 month | ✓ | | |
| 5.3R | IBM | 10 month | | | |
| 5.3L | IBM | 10 month | ✓ | | |
| 5.4R | IBM | 8 month | ✓ | | |
| 5.4L | IBM | 8 month | | | |
| 6.1L | IBM | 4 month | ✓ | | |
| 6.4R | IBM | 8 month | ✓ | | |
| 6.4L | IBM | 8 month | ✓ | | |
| 8.1 | IBM | 4 month | | | |
| 8.9 | IBM | 4 month | ✓ | ✓ | ✓ |
| 8.3 | IBM | 6 month | ✓ | | ✓ |
| 8.5 | IBM | 6 month | | | |
| 8.2 | IBM | 8 month | | | |
| 8.4 | IBM | 8 month | | | |
| 8.12 | IBM | 8 month | | | ✓ |
| 8.7 | IBM | 10 month | ✓ | | ✓ |
| 8.8 | IBM | 10 month | ✓ | ✓ | ✓ |
| 8.11 | IBM | 10 month | ✓ | | ✓ |
| 9.6L | IBM | 4 month | | | |
| 9.5R | IBM | 8 month | | | |
| 9.5L | IBM | 8 month | | | |
| 9.7R | IBM | 8 month | | | |
| 9.7L | IBM | 8 month | | | |

| Xenograft | Group | Treatment | Timepoint | GPSM | HDGF | ACSF |
|---|---|---|---|---|---|---|
| 10.3R | IBM | | 4 month | | | |
| 13.1R | IBM | | 3 month | ✓ | | ✓ |
| 13.1L | IBM | | 3 month | ✓ | ✓ | ✓ |
| 13.2R | IBM | | 4 month | | | |
| 13.2L | IBM | | 4 month | | | |
| 13.4R | IBM | | 6 month | ✓ | | ✓ |
| 13.4L | IBM | | 6 month | ✓ | | |
| 13.3R | IBM | | 8 month | ✓ | | ✓ |
| 13.3L | IBM | | 8 month | | | |
| 13.5R | IBM | | 8 month | | | |
| 13.5L | IBM | | 8 month | | | ✓ |
| 19.3R | IBM | | 4 month | | | |
| 19.4R | IBM | | 4 month | ✓ | | |
| 19.4R | IBM | | 4 month | | | |
| 23.4R | IBM | OKT3 | 2 month | | | |
| 23.4L | IBM | OKT3 | 2 month | | | |
| 23.5R | IBM | Sham | 2 month | ✓ | | ✓ |
| 23.5L | IBM | Sham | 2 month | ✓ | | |
| 23.3R | IBM | Sham | 4 month | ✓ | ✓ | ✓ |
| 23.3L | IBM | Sham | 4 month | ✓ | ✓ | ✓ |
| 23.1 | IBM | OKT3 | 4 month | ✓ | ✓ | ✓ |
| 23.2R | IBM | OKT3 | 4 month | ✓ | ✓ | ✓ |
| 23.2L | IBM | OKT3 | 4 month | ✓ | ✓ | ✓ |
| 26.1 | IBM | Sham | 2 month | ✓ | | |
| 26.4 | IBM | Sham | 2 month | | | ✓ |
| 26.9R | IBM | OKT3 | 2 month | | | |
| 26.9L | IBM | OKT3 | 2 month | | | ✓ |
| 26.2 | IBM | Sham | 4 month | | | |
| 26.5 | IBM | Sham | 4 month | | | ✓ |
| 26.6 | IBM | OKT3 | 4 month | | | |
| 26.7 | IBM | OKT3 | 4 month | | | |
| 26.8 | IBM | OKT3 | 4 month | | | |
| 33.1 | IBM | | 8 month | | | |
| 33.2 | IBM | | 8 month | | | |
| 33.3 | IBM | | 8 month | | | |
| 36.2R | IBM | OKT3 | 4 month | | | |
| 36.2L | IBM | OKT3 | 4 month | | | |
| 36.1 | IBM | Sham | 4 month | | | ✓ |
| 36.3R | IBM | Sham | 4 month | | | |
| 36.3L | IBM | Sham | 4 month | | | |
| 36.4R | IBM | OKT3 | 4 month | | | |
| 36.4L | IBM | OKT3 | 4 month | | | |
| 36.5R | IBM | Sham | 4 month | | | |
| 36.5L | IBM | Sham | 4 month | | | ✓ |
| 36.6R | IBM | OKT3 | 4 month | | | |
| 36.6L | IBM | OKT3 | 4 month | | | |
| 36.7R | IBM | Sham | 4 month | | | |
| 36.7L | IBM | Sham | 4 month | | | |
| 42.4 | IBM | Sham | 4 month | ✓ | ✓ | ✓ |
| 42.5 | IBM | OKT3 | 4 month | ✓ | | ✓ |
| 42.1 | IBM | Sham | 8 month | ✓ | ✓ | ✓ |
| 42.2 | IBM | Sham | 8 month | | | |
| 42.6 | IBM | OKT3 | 8 month | ✓ | ✓ | ✓ |
| 42.7 | IBM | OKT3 | 8 month | | ✓ | |
| 42.8 | IBM | OKT3 | 8 month | | ✓ | ✓ |

**Table S6: Xenograft cryptic exon expression summary.** A summary of the cryptic exon expression for the control and IBM xenograft samples. A checkmark indicates cryptic exon expression for GPSM2 (abbreviated GPSM), HDGFRP2 (HDGF), and ACSF2 (ACSF), and a grey box indicates a cryptic exon target whose expression was not tested. For cases used in OKT3 experiments (23, 26, 36, and 42) Fishers exact test was used to test for significance at each timepoint and no significant differences were found between the groups (2-month, $p > 0.999$; 4-month, $p = 0.3259$; 8-month, $p > 0.999$).
